# Promiscuous esterases counterintuitively are less flexible than specific ones

**DOI:** 10.1101/2020.06.02.129015

**Authors:** Christina Nutschel, Cristina Coscolín, Daniel Mulnaes, Benoit David, Manuel Ferrer, Karl-Erich Jaeger, Holger Gohlke

## Abstract

Understanding mechanisms of promiscuity is increasingly important from a fundamental and application point of view. As to enzyme structural dynamics, more promiscuous enzymes generally have been recognized to also be more flexible. However, examples for the opposite received much less attention. Here, we exploit comprehensive experimental information on the substrate promiscuity of 147 esterases tested against 96 esters together with computationally efficient rigidity analyses to understand the molecular origin of the observed promiscuity range. Unexpectedly, our data reveal that promiscuous esterases are significantly less flexible than specific ones, are significantly more thermostable, and have a significantly increased specific activity. These results may be reconciled with a model according to which structural flexibility in the case of specific esterases serves for conformational proofreading. Our results signify that esterase sequence space can be screened by rigidity analyses for promiscuous esterases as starting points for further exploration in biotechnology and synthetic chemistry.

## 1. Introduction

Enzymes involved in primary metabolism typically exquisitely discriminate against other metabolites. Yet, evolution of specificity is only pushed by nature to the point at which ‘unauthorized’ reactions do not impair the fitness of the organism (1). As a result, the universe of promiscuous activities available in nature has been suggested to be enormous (2, 3). Understanding mechanisms of promiscuity has thus become increasingly important for the fundamental understanding of molecular recognition and how enzyme function has evolved over time(4) but also to optimize enzyme engineering applications (5). A particular challenge in the latter case is the ability to discover a suitable enzyme with ‘sufficient’ promiscuous activity to serve as a starting point for further exploration (1).

Enzyme structural dynamics, besides its role in catalysis (6, 7) and allosteric regulation (8–11), has been recognized as likely the single most important mechanism by which promiscuity can be achieved (5). Prominent examples are human cytochrome P450 (CYP) enzymes, for which crystallographic studies and molecular simulations demonstrated that more promiscuous CYPs show larger structural plasticity and mobility (12–14), or TEM-1 β-lactamase and a resurrected progenitor, for which molecular simulations show that the pocket of the ancestral, and more promiscuous, enzyme fluctuates to a greater extent (15). However, examples for the opposite, i.e., conformational changes selected in evolution such that they enhance specificity in molecular recognition (16), have received much less attention in the context of enzyme promiscuity.

A clear limitation for scrutinizing the link between enzyme structural dynamics and substrate promiscuity is the general lack of large-scale data on one enzyme (super)family tested against a multitude of ligands (17) (cf. ref. (1) for notable exceptions). Likewise, acquiring information on enzyme dynamics at the atomistic level by experimental techniques or classical molecular dynamics (MD) simulations is burdensome. Here, we exploit comprehensive experimental information on the substrate promiscuity (18) of esterases (abbreviated EHs, for “Ester Hydrolases”) (19) together with computationally efficient rigidity analyses (20–23) of comparative models of EHs to understand the molecular origin of the observed promiscuity range. Enzyme rigidity, or its opposite flexibility, are static properties that denote the *impossibility*, or *possibility*, *of motions* in an enzyme under force, without giving information about directions and magnitudes of movements (23). Enzyme flexibility, thus, should not be confused with enzyme mobility, which describes *actual motions* in an enzyme. Rigidity analysis results do not rely on the correct description of the time-dependency of processes (23), which makes them valuable in cases where timescales over multiple orders of magnitude may govern such processes, like in enzyme dynamics (6, 7).

In recent years, EHs have obtained much attention in basic research and industrial applications (24). EHs are widely distributed in nature within microbial communities (at least one EH is found in each bacterial genome), they have been extensively examined with state-of-the-art (meta)genomics techniques and investigated by functional screenings compared to many other classes of enzymes. They also possess outstanding properties in terms of stability, reactivity, and scalability that make them appropriate biocatalysts to improve competitiveness, innovation capacity, and sustainability in a modern circular bio-economy (25). Recently, a large-scale study on substrate promiscuity (*P*_EH_, which denotes the number of esters hydrolyzed by an EH) of 147 phylogenetically, environmentally, and structurally diverse microbial EHs was described by Martínez-Martínez *et al.* (19), in which all EHs were functionally assessed against a customized library of 96 esters. As to mechanistic understanding, the authors related *P*_EH_ to a structural parameter, the active site effective volume. However, the impact of enzyme flexibility on *P*_EH_ was not assessed.

In our study, we thus ask the following questions: I) What is the relation between *P*_EH_ and EH flexibility? II) Does this relation hold if experimentally determined EH thermostabilities are used as proxies for enzyme flexibility? III) What is the relation between *P*_EH_ and EHs’ specific activities? IV) Is there a preference of promiscuous or specific EHs for a particular type of esters. V) Can this preference be understood with respect to EHs flexibilities?

## 2. Materials and Methods

### 2.1. Definition of data sets

The present study builds on the study from Martínez-Martínez *et al.* (19). In order to assess *P*_EH_, the authors experimentally investigated 147 phylogenetically, environmentally, and structurally diverse microbial EHs (termed *experimental data set*) against a customized library of 96 different esters. Two commercial lipases, which have found wide biotechnological applications, CalA and CalB from *Pseudozyma aphidis* (formerly *Candida antarctica*), were included for comparison. For details on determining and classifying *P*_EH_, see **Supplemental Materials and Methods**. To validate that *P*_EH_ defines promiscuity of EHs in a quantitative manner, *k*_cat_ and *K*_m_ values were determined for ten expressed and purified EHs covering the entire *P*_EH_ range (see **section 2.9**) and a promiscuity index *I* (**see Supplemental Materials and Methods**, **Eq. S4**) computed and compared to *P*_EH_. Finally, the similarity of the ester substrates was assessed by the maximum pairwise Tanimoto-Combo similarity score *δ*_*ij*_ for compound *i* versus *j*, accounting for shape and chemical complementary between 3D structures, and the mean maximum pairwise Tanimoto-Combo similarity score *δ*_*ij*_ of a substrate *i* to all other substrates in the data set (see **Supplemental Methods and Materials**).

As our computational approach involves extensive molecular dynamics (MD) simulations for generating conformational ensembles **(see section 2.3)**, we selected 35 EHs from the *volume data set* (termed *flexibility data set*) for comparative modeling **(see section 2.2)**. The criteria for choosing EHs of the *flexibility data set* are explained in **section 3.1**.

### 2.2. Comparative modelling and validations of the *flexibility data set*

Comparative models of the *flexibility data set* **(see section 2.1)** were generated using our in-house structure prediction meta-tool TopModel (26) that has been successfully applied in previous studies (27–30). TopModel uses multiple state-of-the-art threading and sequence/structure alignment tools to generate a large ensemble of models from different pairwise and multiple alignments of the top five highest ranked template structures. The TopModel software is available at https://cpclab.uni-duesseldorf.de/index.php/Software.

The quality of the homology models was assessed by our meta Model Quality Assessment Program (meta-MQAP) TopScore (31). TopScore uses deep neural networks (DNN) to combine scores from 15 different primary MQAP to predict accurate residue-wise and whole-protein error estimates. For details on model quality assessment by TopScore and validation, see **Supplemental Materials and Methods**.

To test whether CARs of the homology models are accessible for substrates, we applied the CAVER 3.0.3 PyMOL Plugin (32). Starting points for the computations were defined based on the Cartesian coordinates of the CARs’ center of mass (COM). Default values were used for the probe radius (0.9 Å), shell radius (3.0 Å), and shell depth (4.0 Å).

### 2.3. Generation of structural ensembles

Structural ensembles of EHs were generated by all-atom MD simulations of in total 5 μs simulation time per EH. For details on starting structure preparation, parametrization, and equilibration see **Supplemental Materials and Methods**.

All minimization, equilibration, and production simulations were performed with the *pmemd.cuda* module (33) of Amber19 (34). During production simulations, we set the time step for the integration of Newton’s equation of motion to 4 fs following the hydrogen mass repartitioning strategy (35). Coordinates were stored into a trajectory file every 200 ps. This resulted in 5000 configurations for each production run that were considered for subsequent analyses.

### 2.4. Constraint Network Analysis

The flexibility analyses were performed with the Constraint Network Analysis (CNA) software package (version 3.0) (20–23). CNA functions as front- and back-end to the graph theory-based software Floppy Inclusions and Rigid Substructure Topography (FIRST) (36). Applying CNA to biomolecules aims at identifying their composition of rigid clusters and flexible regions, which can aid in the understanding of biomolecular structure, stability, and function (21–23). As the mechanical heterogeneity of biomolecular structures is intimately linked to their diverse biological functions, biomolecules generally show a hierarchy of rigidity and flexibility (20). In CNA, biomolecules are modeled as constraint networks in a *body-and-bar* representation, which has been described in detail by Hesphenheide *et al*. (37). A fast combinatorial algorithm, the *pebble game*, counts the bond rotational degrees of freedom and floppy modes (internal, independent degrees of freedom) in the constraint network (38). In order to monitor the hierarchy of rigidity and flexibility of biomolecules, CNA performs thermal unfolding simulations by consecutively removing non-covalent constraints (hydrogen bonds, including salt bridges) from a network in increasing order of their strength (39–41). For details on thermal unfolding simulations, see **Supplemental Materials and Methods**. To improve the robustness and investigate the statistical uncertainty, we carried out CNA on ensembles of network topologies (ENT^MD^) generated from MD trajectories **(see section 2.3)** (42).

The CNA software is available under academic license at https://cpclab.uni-duesseldorf.de/index.php/Software and the CNA web server is accessible at https://cpclab.uni-duesseldorf.de/cna.

### 2.5. Local and global indices

From the thermal unfolding simulations, CNA computes a comprehensive set of indices to quantify biologically relevant characteristics of the protein’s stability. *Global* indices are used for determining the rigidity and flexibility at a macroscopic level; *local* indices determine the rigidity and flexibility at a microscopic level of bonds (43). The cluster configuration entropy *H*_type2_ is a *global* index that has been introduced by Radestock and Gohlke (20). As done previously, we applied *H*_type2_ as a measure for global structural stability of proteins (20, 41, 44–48). The stability map *rc*_*ij*_ is a *local* index that has been introduced by Radestock and Gohlke (20). We applied *rc*_*ij*_ as a measure for local structural stability of proteins in previous studies (45, 47, 48). For details on both indices, see **Supplemental Materials and Methods**.

### 2.6. Root mean square fluctuations

The per-residue root-mean-square fluctuations were calculated for each EH (*RMSF*_EH_) and for its CARS (*RMSF*_CAR_) based on the MD trajectories **(see section 2.3)**. Prior to the calculations, the structures of each trajectory were superimposed onto the average structure using the 90% least mobile residues of the respective EHs (49).

### 2.7. Torsion angles

For each of the 96 esters, the number of freely rotatable bonds (torsion angles, TA) was calculated based on the SMILES codes provided by Martínez-Martínez *et al.* (19).

To compare how many esters with a specific TA are hydrolyzed by each EH, we calculated the normalized proportion of ester hydrolysis with a specific TA (*Norm*_ester_(TA)) as the number of hydrolyzed esters with a specific TA (*Ester*_hydrolysed_ (TA)) divided by the total number of esters in the data set with this specific TA (*Ester*^library^ (TA)) (Eq. 1).

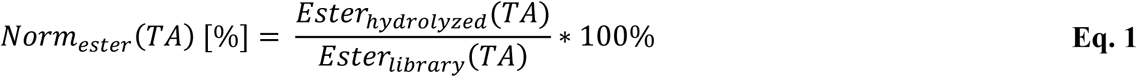

### 2.8. Circular dichroism spectroscopy

Eleven serine ester hydrolases, EH_1_ (Protein data Bank acc. nr. 5JD4), EH_2_ (GenBank acc. nr. KY483643), EH_3_ (GenBank acc. nr. KY483645), EH_4_ (GenBank acc. nr. KR107250), EH_6_ (GenBank acc. nr. KP347751), EH_8_ (GenBank acc. nr. WP_011587341.1), EH_9_ (GenBank acc. nr. KY483648), EH_16_ (GenBank acc. nr. KP347759), EH_21_ (GenBank acc. nr. KP347760), EH_37_ (GenBank acc. nr. KR107248), EH_43_ (GenBank acc. nr. KP347758) from metagenomic origin, were used in the present study to perform CD determinations. The vector pET46 Ek/LIC and the host *Escherichia coli* MC1061 were the source of the His_6_-tag EH_1_, EH_4_, EH_8_, and EH_37_, the vector pBXNH3 and the host *E. coli* MC1061 were the source of the His_6_-tag EH_2_, EH_3_, and EH_9_, and the vector p15Tv-L and the host E. coli BL21(DE3) were the source of EH_6_, EH_16_, EH_21_, and EH_43_. Prior to analyses, the soluble His-tagged proteins were produced and purified after binding to a Ni-NTA His-Bind resin (Sigma-Aldrich, MO, US) as described by Martínez-Martínez (19). Purity was assessed as >98% using SDS-PAGE analysis in a Mini PROTEAN electrophoresis system (Bio-Rad, Madrid, Spain) and subsequent staining with Coomassie Brilliant Blue. A total of about 10-20 mg total purified recombinant protein was obtained on average from 1-liter culture. All proteins were stored at −20°C at a concentration of 10 mg/ml in 40 mM (4-(2-hydroxyethyl)-1-piperazineethanesulfonic acid (HEPES) buffer (pH 7.0), until used. Circular dichroism (CD) spectra were acquired between 190 and 270 nm with a Jasco J-720 spectropolarimeter equipped with a Peltier temperature controller, employing a 0.1 mm cell at 25°C. Spectra were analyzed, and denaturation temperatures were determined at 220 nm between 10 and 85°C at a rate of 30°C per hour, in 40 mM (4-(2-hydroxyethyl)-1-piperazineethanesulfonic acid (HEPES) buffer, pH 7.0. A protein concentration of 1.0 mg ml^−1^ was used. Denaturation temperatures were calculated by fitting the ellipticity (mdeg) at 220 nm at each of the different temperatures using a 5-parameters sigmoid fit with Sigma Plot 14.0.

### 2.9. Kinetic parameter determination

Ten serine ester hydrolases, EH_1_ (Protein data Bank acc. nr. 5JD4), EH_3_ (GenBank acc. nr. KY483645), EH_5_ (GenBank acc. nr. KR107271), EH_7_ (GenBank acc. nr. KY483644), EH_12_ (GenBank acc. nr. KR107263), EH_17_ (GenBank acc. nr. KR107278), EH_37_ (GenBank acc. nr. KR107248), EH_102_ (Protein data Bank acc. nr. 5JD3), EH_114_ (GenBank acc. nr. KR107274) and EH_127_ (GenBank acc. nr. KR107253) from metagenomic origin, were used in the present study to perform kinetic determinations (*k*_cat_ and *K*_m_). The vector pET46 Ek/LIC and the host *Escherichia coli* MC1061 were the source of the His_6_-tag EH_1_, EH_5_, EH_12_, EH_17_, EH_37_, EH_102_, EH_114_, and EH_127_, and the vector pBXNH3 and the host *E. coli* MC1061 was the source of the His^6^-tag EH_3_. The soluble His-tagged proteins was produced and purified as described by Martínez-Martínez (19). For details see above.

The kinetic parameters were calculated at 550 nm using a continuous pH indicator (Phenol red; ε550 nm = 8450 M^−1^ cm^−1^) assay at 550 nm in 384-well plates as previously described (Alonso et al, 2020). Briefly, to 40 µl of 5 mM 4-(2-hydroxyethyl)-1-piperazinepropanesulfonic acid (EPPS) buffer (pH 8.0) 2 µl of a stock ester solution was added to achieve the desired concentration of each ester. Finally, 2 µl of stock protein solution was immediately added to each well, to achieve the desired protein concentration, using an Eppendorf Repeater M4 pipette (Eppendorf, Hamburg, Germany). The total reaction volume was 44 µl. Ester hydrolysis was measured at 30°C in a Synergy HT Multi-Mode Microplate Reader in continuous mode at 550 nm over 24 h. One unit (U) of enzyme activity was defined as the amount of free enzyme or enzyme bound to the carrier required to transform 1 µmol of substrate in 1 min under the assay conditions using the reported extinction coefficient (Phenol red at 550 nm = 8450 M^−1^ cm^−1^). For *K*_*m*_ determination - [protein]: 4.5 μg ml^−1^; [ester]: 0-100 mM; reaction volume: 44 μl; T: 30°C; pH: 8.0. For *k*_cat_ determination - [protein]: 0-270 μg ml^−1^; [ester]: 100 mM; reaction volume: 44 μl; T: 30°C; pH: 8.0. All values, in triplicates, were corrected for non-enzymatic transformation. Kinetic parameters were calculated by fitting the data fit with Sigma Plot 14.0.

## 3. Results

### 3.1. Definition of data sets

To understand the structural origin of and develop a method to predict *P*_EH_, the present study builds on large-scale data from Martínez-Martínez *et al.* (19). The authors experimentally investigated *P*_EH_ of 147 EHs (termed *experimental data set*) **(see section 2.1)**. In doing so, compromises needed to be made regarding the measurement of catalytic activity, i.e., specific activity was measured using enzymes expressed in *E. coli* without subsequent purification, only a single concentration of wet cells expressing enzymes (0.4 mg per ester) was used to measure activity, and the substrates were tested at a single concentration of circa 7 mM on average (19). To validate that the *P*_EH_ derived from the measured activities define promiscuity of the enzymes in a quantitative manner, *k*_cat_ and *K*_m_ values were determined now for ten expressed and purified EH covering the entire *P*_EH_ range **(see section 2.9)**. *k*cat/*K*m ideally serves as the kinetic parameter in enzyme promiscuity studies for comparison (50, 51). From *k*_cat_/*K*_m_ values of an enzyme towards a defined set of substrates, a quantitative index of promiscuity *I* **(Eq. S4)** can be calculated based on information entropy (51). *I* yields a very good and significant (*R*^2^ = 0.79, *p* = 0.0003) correlation with *P*_EH_, indicating that *P*_EH_ relates to EH promiscuity in a quantitative manner **(Figure S1)**, although the range of *I* suggests that large *P*_EH_ may still be associated with moderate promiscuity. Note that, although the promiscuity index is a functional parameter that is defined for a specified set of substrates, promiscuity indices for different enzymes are quantitatively comparable if they have been calculated using the same substrate set (51). Furthermore, for the ten EH and using the colorimetric assay herein used, *k*_cat_/*K*_m_ can be measured with a standard error of the mean as low as 0.05 min^−1^ mM^−1^, which corresponds to *k*_cat_ and *K*_m_ fitting values 2-fold above the background signals under assay conditions for each of the enzymes and esters. When *k*_cat_/*K*_m_ > 0.05 min-1 mM-1 is used as a criterion to define that a substrate is hydrolyzed, the resulting number of substrates for the ten EHs yields an excellent and significant (*R*^2^ = 0.99, *p* < 10^−4^) correlation with *P*_EH_ **(Figure S2)**, again indicating that *P*_EH_ relates to EH promiscuity in a quantitative manner. Finally, esters that are chemically similar to each other are expected to be metabolized similarly by an EH; such correlations in the substrate set would reduce the effective EH promiscuity. Therefore, similarity of the substrates was assessed by the maximum pairwise Tanimoto-Combo similarity score *δ*_*ij*_ for compound *i* versus *j*, which is bounded between 0 for dissimilar compounds and 2 for identical ones, and the mean maximum pairwise Tanimoto-Combo similarity score *δ*_*i*_ of a substrate *i* to all other substrates in the data set; the Tanimoto-Combo similarity score accounts for shape and chemical complementary between 3D structures as determined by the Rapid Overlay of Chemical Structures approach (52). Complete linkage clustering on the pairwise distance matrix calculated for all 96 compounds from *δ*_*ij*_ yielded 20 clusters at a distance of 1.0 (**Figure S3**), which is equivalent to *δ*_*ij*_ = 1.0, indicating that on average less than five esters share a similarity that is half-way between dissimilar and identical. The negatively skewed histogram of *δ*_*i*_ furthermore shows that *δ*_*i*_ peaks at 1.0 and is below 1.2 (**Figure S4**), indicating that an ester generally shares a similarity to all other esters that is half-way between dissimilar and identical, or worse.

Additionally, Martínez-Martínez *et al.* ranked (classified) *P*_EH_ of 96 EHs (termed *volume data set*) based on the active site effective volume **(see section 2.1) (Eq. S1)** (19), which will be used here as a reference to compare the power of *P*_EH_ predictions. As our computational approach involves extensive MD simulations for generating conformational ensembles **(see section 2.3)**, we selected 35 EHs from the *volume data set* based on the following criteria; they constitute the *flexibility data set*. I) The data set contains eleven EHs with known crystal structures (including the commercial EHs CalA and CalB) (Figure 1A, **Table S1)** and 24 EHs for which no experimental structure is available but for which comparative models can be generated **(see section 3.2) (Figure 1A, **Table S2)**. That way, we can probe to what extent the source of structural information influences the outcome of our results. II) The chosen EHs of the data set show high diversities as to *P*_EH_ and association to esterase families (*F*_EH_, as defined by Arpigny and Jaeger (53)), similar to those found for the *volume data set* **(Figures S1 and S2, Tables S3 and S4)**. This resulted in *P*_EH_ ranging from 4 to 72 (Figure 1A, Tables S1 and S2)**. In the following, we consider *P*_EH_ as *low* if the EH hydrolyzes ≤ 9 esters (11% of the data set), as *moderate* if the EH hydrolyzes between 10 and 29 esters (49%), and as *high* if the EH hydrolyzes ≥ 30 esters (40%) **(Figure S5, Table S3)**. The data set covers eleven *F*EH of which FIV (44% of the data set) and FV (21%) are the best represented ones **(Figure S6, Table S4)**. This reflects the proportion of their presence in the *volume data set*. III) Only EHs with amino acid sequence identities ≥ 25% in comparison to an existing crystal structure were considered **(see section 2.1)** in order to ensure a sufficient quality of generated comparative models.

**Figure 1:**
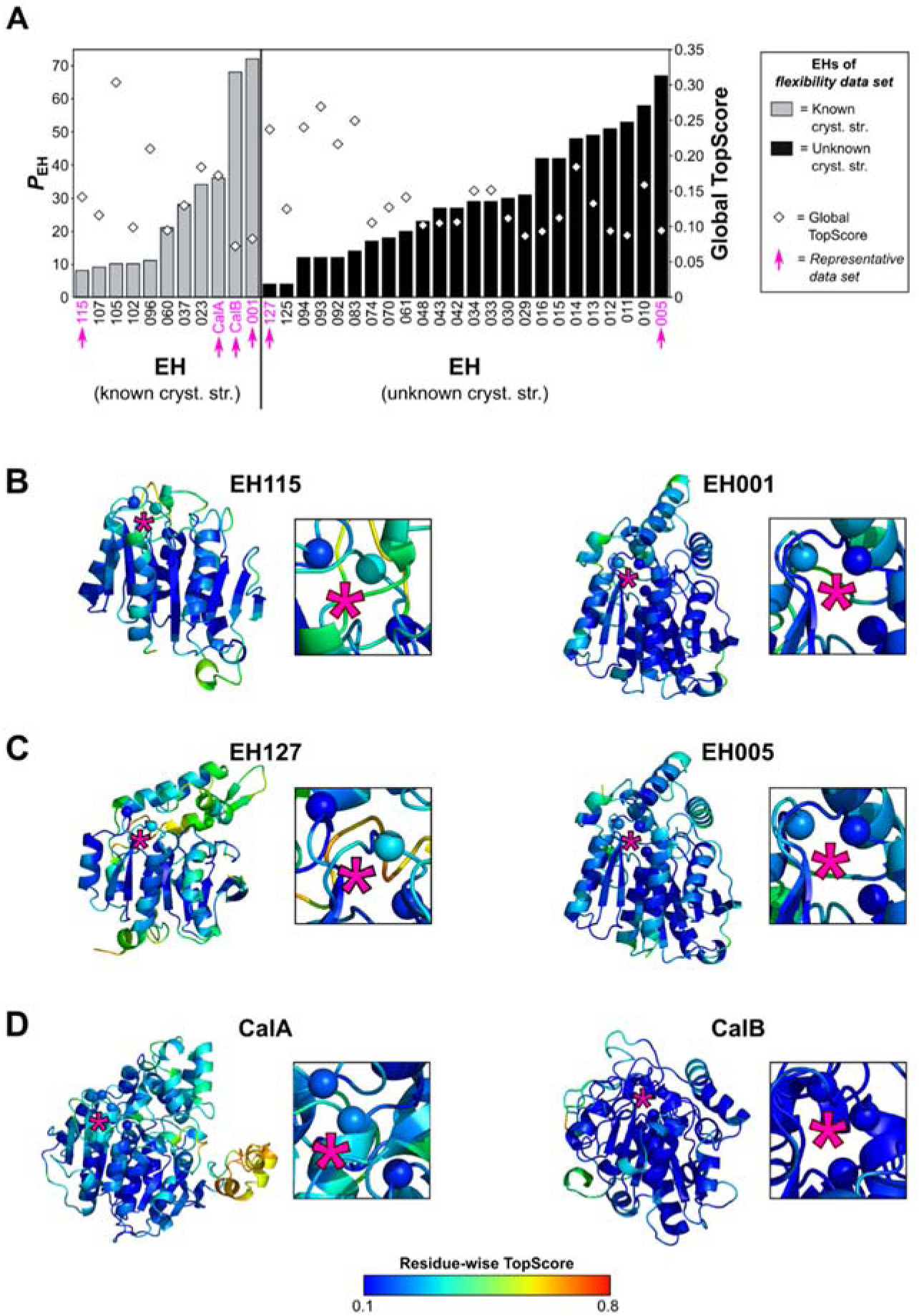
Comparative modeling of EHs. **(A)** Based on sequence data provided by a large-scale study from Martínez-Martínez *et al.* (19), comparative models were generated for 35 EHs with known (left, 11 EHs) and unknown (right, 24 EHs) crystal structures using TopModel (26). These EHs constitute the *flexibility data set*. The EHs vary in *P*_EH_ (left ordinate, bars) and global TopScores (right ordinate, diamonds). Six EHs were selected as representatives of the *flexibility data set* (termed *representative data set*) as indicated by magenta arrows. The quality of the comparative models of **(B)** EHs with known crystal structures and lowest (EH115) or highest *P*_EH_ (EH001), **(C)** EHs with unknown crystal structures and lowest (EH127) or highest *P*_EH_ (EH005), and **(D)** commercial EHs with highest (CalA) or lowest *P*_EH_ (CalB) was evaluated by TopScore (31). For each comparative model the residue-wise TopScore is shown: A good (bad) homology model shows a low (high) residue-wise TopScore (see color scale at the bottom). Insets depict CARs (spheres) within an EH. For clarity the position of CARs is indicated by magenta stars.

Finally, in order to uniformly depict the results across the present study, six EHs were selected as representatives of the *flexibility data set* based on *P*_EH_ (termed *representative data set*): EHs with the lowest (EH115) or highest *P*_EH_ (EH001) and known crystal structures, EHs with the lowest (EH127) or the highest *P*_EH_ (EH005) and unknown crystal structures, and commercial EHs with the lowest (CalA) or highest *P*_EH_ (CalB) (Figure 1A-D, **Tables S1 and S2)**.

### 3.2. Comparative models of EHs generated by TopModel show an overall and residue-wise good quality

To generate structural models of EHs as starting points for our investigations, we performed template-based modeling of the *flexibility data set* using TopModel (26) **(see section 2.2)**. In doing so, we also generated comparative models of the eleven EHs for which crystal structures are available. These structural models will be used to judge the quality of the comparative modeling.

The quality of the comparative models of the *flexibility data set* were assessed with TopScore (31), a meta Model Quality Assessment Program (meta-MQAP) **(see section 2.2)**. For the eleven Es with known crystal structure, the global TopScores range from 0.074 to 0.305 (Figure 1A, **Table S1)**. As the global TopScore describes whole-protein error estimates, this shows that the structures contain between 7.4 and 30.5% error. Notably, the global TopScores well and significantly correlate (*R*^2^ = 0.61, *p* = 0.004) with values of 1 – lDDT computed from comparisons of the comparative models of EHs with known crystal structure against these experimental reference structures, indicating that global TopScores are well suited to assess the model quality of EHs (**Figure S7, Table S5**). The global TopScore values of the comparative models of the other 24 EHs range from 0.087 to 0.269 (Figure 1A, **Table S2)**, indicating that these models are of equal quality than the ones for EHs with known crystal structure. The TopScore values of the *representative data set* lie in a comparable range (Figure 1A, **Tables S1 and S2)**. Moreover, the comparative models of the *flexibility data set* show low residue-wise TopScore values (31), indicating that all parts of a model are of good quality. We illustrate this for the residue-wise TopScore of the comparative models of the *representative data set* (Figures 1B-D). This also applies to structural regions around CARs (Figures 1B-D). That way, it was possible to confirm CARs in models of EHs with known crystal structures and to unambiguously identify CARs in models of EHs with unknown crystal structures (Figures 1B-D, **Tables S1 and S2)**.

Additionally, we validated that CARs in all models are accessible for substrates according to CAVER results (32) **(see section 2.2)**, i.e., that all models are in an open conformation: CARs are either located on the protein surface or are buried and connected with the surface by tunnels. We illustrate this for the comparative models of the *representative data set* **(Figure S8)**.

To conclude, comparative models were generated for 35 EHs of the *flexibility data set* using TopModel. The models showed both an overall and residue-wise good structural quality. Additionally, we validated that CARs in all models are accessible for substrates.

### 3.3. Promiscuous EHs are globally less flexible

Previous studies indicated that enzyme flexibility influences the substrate promiscuity of enzymes (12–14). For gaining insights into how the flexibility of EHs is linked to *P*_EH_, we applied CNA (21, 23), a rigidity theory-based approach to analyze biomolecular statics (21–23), to the *flexibility data set* **(see sections 2.4)**. To improve the robustness and investigate the statistical uncertainty, for each of the comparative models we carried out CNA on ensembles of network topologies (ENT^MD^) generated from five MD trajectories of 1 μs length each (44) **(see sections 2.3 and 2.4)**. In order to investigate if the global flexibility of the EHs influences *P*_EH_, we predicted *T*_p_, the phase transition temperature previously applied as a measure of structural stability of a protein (20, 41, 44–48), for each EH **(see section 2.5)**. *T*_p_ was averaged over five ensembles **(see sections 2.3 and 2.4)**, resulting in all but one case in SEM < 1.87 K (Figure 2A, **Tables S1 and S2)**.

**Figure 2:**
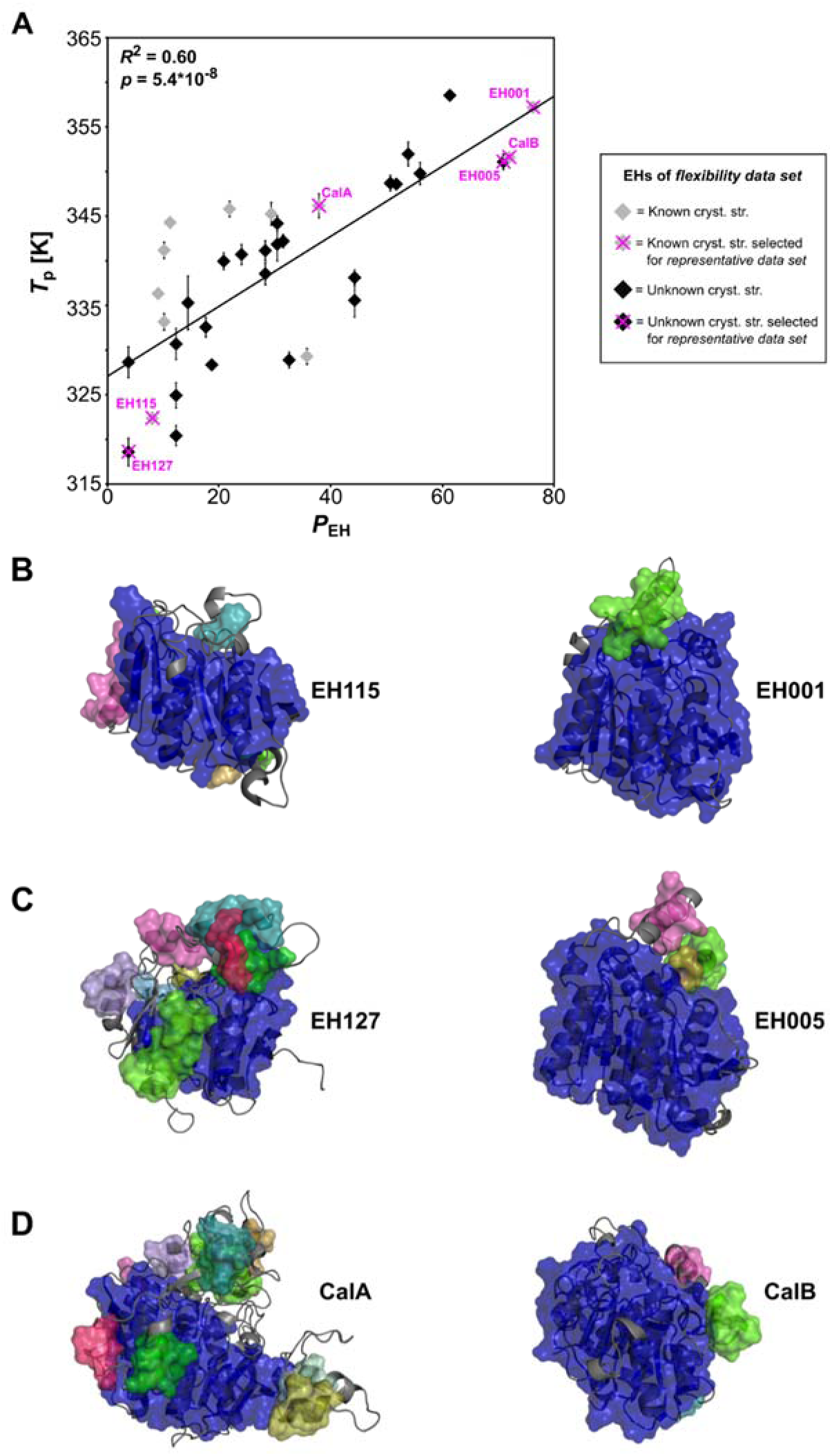
Correlation of *T*_p_ *versus P*_EH_. **(A)** Correlation between predicted *T*_p_ based on the global index *H*type2 and *P*_EH_ for the *flexibility data set*. Data points colored grey (black) represent comparative models of EHs with (un)known crystal structures. The *representative data set* is indicated by magenta crosses. Error bars show the SEM over five independent MD simulations of 1 µs length each. Rigid cluster decomposition at 332 K during the thermal unfolding simulation of **(B)** EHs with known crystal structures and lowest (EH115) or highest *P*_EH_ (EH001), **(C)** EHs with unknown crystal structures and lowest (EH127) or highest *P*_EH_ (EH005), and **(D)** commercial EHs with lowest (CalA) or highest *P*_EH_ (CalB). Rigid clusters are represented as uniformly colored blue, green, pink, cyan, and magenta bodies in the descending order of their sizes.

*T*_p_ and *P*_EH_ of the *flexibility data set* are well and significantly correlated (*R*^2^ = 0.60, *p* = 5.4^*^10^−8^) (Figure 2A). To validate the consistency of our approach, we considered EHs with known or unknown crystal structures separately. In both cases, good and significant correlations between *T*_p_ and *P*_EH_ were revealed (known crystal structures: *R*^2^ = 0.48, *p* = 0.019; unknown crystal structures: *R*^2^ = 0.73, *p* = 1.1^*^10^−7^), lending support to the quality of comparative models predicted with TopModel and indicating that future predictions on EHs with unknown experimental structures should be promising. Notably, EHs with high *P*_EH_ have a high *T*_p_ and *vice versa*, i.e., promiscuous EHs are globally less flexible. Exemplarily, this is depicted for EHs of the *representative dataset* with known crystal structures and lowest (EH115) or highest *P*_EH_ (EH001), which showed *T*_p_ of 322.3 K and 357.2 K, with unknown crystal structures and lowest (EH127) or highest *P*_EH_ (EH005), which showed *T*_p_ of 318.6 K and 351.1 K, and CalA and CalB, which showed *T*_p_ of 346.2 K and 351.6 K (Figure 2A, **Tables S1 and S2)**. The differences in global structural stability of these EHs are illustrated by the rigid cluster decomposition at 332 K during the thermal unfolding simulations (Figure 2B-D): promiscuous EHs are globally more structurally stable at the elevated temperature as indicated by fewer, but larger, rigid clusters.

The EH flexibility analyzed so far is a static property and describes the potential of motions in a biomolecule (23). Yet, direct information on mobility within EHs is available from the ensembles generated by MD simulations. We thus computed exemplarily *RMSF*EH, a measure for protein mobility **(see section 2.6)**, across the ensembles of EHs from the *representative data set*. *RMSF*EH, averaged over all residues and all five MD trajectories, and *P*_EH_ do not yield a significant correlation (*p* = 0.13) **(Figure S9A, Table S6)**, in contrast to *T*_p_ and *P*_EH_ (*R*^2^ = 0.93, *p* = 1.8^*^10^−3^) **(Figure S9B, Table S6)**. Still, as promiscuous EHs are globally less mobile, the same trend is obtained as in the case of the flexibility analysis.

To conclude, a good and significant correlation between *T*_p_ and *P*_EH_ was found for the *flexibility data set* (*R*^2^ = 0.60, *p* = 5.4^*^10^−8^). These findings demonstrate that promiscuous EHs are globally less flexible. *RMSF*EH is less predictive for *P*_EH_, although again promiscuous EHs are characterized by a lower global mobility, mutually confirming either result.

### 3.4. Promiscuous EHs are more thermostable

Previous studies indicated that thermodynamically more thermostable proteins frequently have a higher structural stability (45, 48). In order to investigate if promiscuous EHs, which were predicted to be less flexible **(see section 3.3)**, are also more thermostable, CD spectroscopy was applied to determine the melting temperature *T*_d_ of the EHs **(see section 2.8)**. Note that only if the unfolding of a protein is reversible, CD spectroscopy provides true thermodynamic properties (54). However, even if the unfolding is irreversible, because the protein aggregates at high temperatures, the method can still give information about relative stabilities (54). Hence, to reduce the potential impact of different aggregation kinetics of structurally different proteins, we applied CD spectroscopy to one *F*EH family only. In particular, we used *F*_IV_ because it is the largest *F*EH **(Table S7)**.

Exemplarily, a CD spectrum for *T*_d_ determination is shown for EH001 (Figure 3A); for each EH, *T*_d_ determination was performed in triplicates with STD < 0.62 K. *T*_d_ and *P*_EH_ yield a fair and significant correlation (*R*^2^ = 0.32, *p* = 0.033, Figure 3B; a similarly fair and significant correlation is found if the data point with highest *T*_d_ is omitted (*R*^2^ = 0.45, *p* = 0.016)).

**Figure 3:**
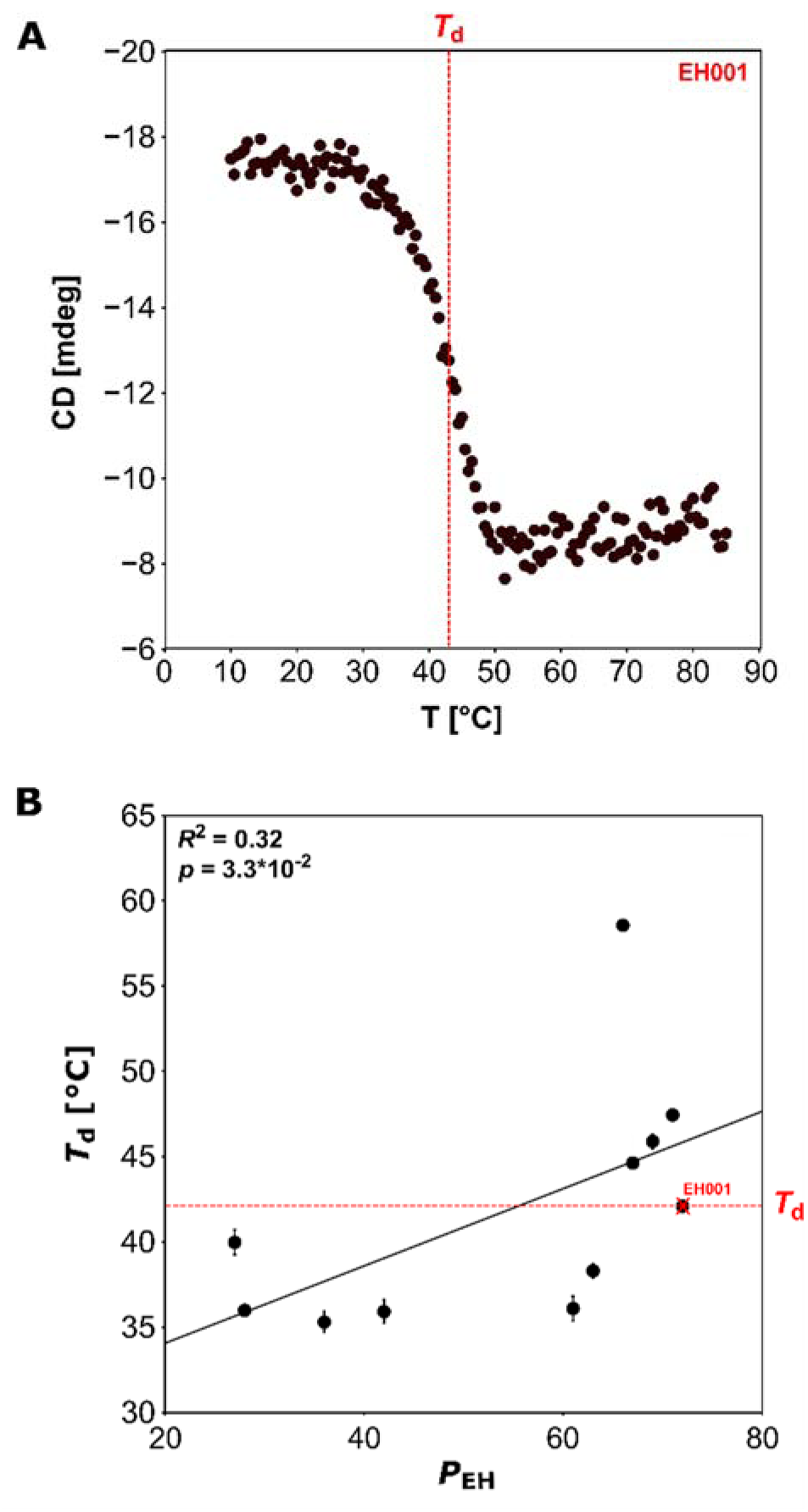
Determination of *T*_d_ via CD spectroscopy. **(A)** Exemplary CD spectrum of EH001. The ellipticity changes in mdeg at 220 nm was plotted against the temperature, resulting in a sigmoidal curve. The inflection point was used to obtain the *T*_d_ value (dotted line). **(B)** Correlation between *T*_d_ and *P*_EH_ for 12 EHs of FIV.

To conclude, promiscuous EHs are not only globally less flexible but also more thermostable.

### 3.5. Promiscuous EHs have less flexible catalytically active residues

The good correlation of *P*_EH_ and *T*_p_ encouraged us to investigate if local flexibility characteristics of CARs will provide an even better predictor of EH promiscuity. We thus computed *Flex*CAR for the *flexibility data set*, i.e., the stability of rigid contacts between CARs and other residues that are at most 5 Å apart from each other, based on the *local* index *rc*_*ij,neighbor*_ **(see section 2.5)**. For each EH, *Flex*CAR was averaged over five ensembles **(see sections 2.3 and 2.4)**, resulting in SEM < 0.06 kcal mol^−1^ **(Figure S10A, Tables 1 and 2)**.

*Flex*_CAR_ and *P*_EH_ of the *flexibility data set* yield a good and significant correlation (*R*^2^ = 0.51, *p* = 1.7^*^10^−6^) **(Figure S10A)**. To validate again the consistency of our approach, we considered EHs with known and unknown crystal structures separately. In both cases, good and significant correlations between *Flex*CAR and *P*_EH_ were found (known crystal structures: *R*^2^ = 0.63, *p* = 3.7^*^10^−3^; unknown crystal structures: *R*^2^ = 0.47, *p* = 2.2^*^10^−4^), again lending support to the quality of comparative models predicted with TopModel. Hence, EHs with high *P*_EH_ have low *Flex*CAR and *vice versa*, i.e., promiscuous EHs have less flexible CARs. Exemplarily, this is detailed for EHs of the *representative dataset* with known crystal structures and lowest (EH115) or highest *P*_EH_ (EH001), which showed *Flex*CAR of −0.74 kcal mol^−1^ and −1.91 kcal mol^−1^, with unknown crystal structures and lowest (EH127) or highest *P*_EH_ (EH005), which showed *Flex*CAR of −1.10 kcal mol^−1^ and −1.86 kcal mol^−1^, and CalA and CalB, which showed *Flex*CAR of −1.31 kcal mol^−1^ and −1.95 kcal mol^−1^ **(Figure S10A, Tables S1 and S2)**. The differences in local structural stability of these EHs are illustrated by rigid contacts between CARs and other residues that are at most 5 Å apart from each other **(Figure S10B-D)**: promiscuous EHs are locally more structurally stable as indicated by more stable rigid contacts.

Finally, we exemplarily computed *RMSF*CAR, a measure for the mobility of a protein’s CARs **(see section 2.6)**, across the ensembles of EHs from the *representative data set*. Averaged *RMSF*CAR and *P*_EH_ correlate worse (*R*^2^ = 0.74, *p* = 0.029) **(Figure S11A, Table S6)** than *Flex*CAR and *P*_EH_ (*R*^2^ = 0.92, *p* = 2.4^*^10^−3^) **(Figure S11B, Table S6)**, paralleling the above results for the global measures. Still, again, as promiscuous EHs have less mobile CARs, the same trend is obtained as in the case of the flexibility analysis.

To conclude, a good and significant correlation between *Flex*CAR and *P*_EH_ was found for the *flexibility data set* (*R*^2^ = 0.51, *p* = 1.7^*^10^−6^). Hence, promiscuous EHs have less flexible CARs. *RMSF*CAR is less predictive for *P*_EH_, although again promiscuous EHs are characterized by less mobile CARs, mutually confirming either result.

### 3.6. Promiscuous EHs have an increased specific activity

In the study by Martínez-Martínez *et al.* (19), the *experimental data set* was screened against 96 esters in a kinetic pH indicator assay **(see section 2.1)**. Besides the average specific activity *Act*_average_ given in U / (g wet cells), also the average maximum specific activity *Act*_max_ was determined. Motivated by the reactivity-selectivity principle (RSP) initially introduced for organic chemistry reactions (55), which states that a more reactive chemical compound is less selective in chemical reactions, we intended to probe if *P*_EH_ is related to *Act*_*max*_. For this, we established an approximate linear free-energy relationship (LFER) (56) by relating log(*Act*_max_) and log(*P*_EH_) **(Figure S12, Table S8)**. In this analysis, the CalA and CalB preparations were excluded because *Act*_max_ was given in U / (g total protein) there.

Log(*Act*_max_) and log(*P*_EH_) of the *experimental data set* yield a good and significant correlation (*R*^2^ = 0.50, *p* = 4.6^*^10^−23^) **(Figure S12A)**. Likewise, log(*Act*_max_) and log(*P*_EH_) of the *flexibility data set* yield a fair and significant correlation (*R*^2^ = 0.22, *p* = 0.6^*^10^−2^) **(Figure S12B)**. To validate whether the same trend emerges for EHs with known and unknown crystal structures, we considered both types of EHs separately. In both cases, fair and significant correlations between log(*Act*_max_) and log(*P*_EH_) were found (known crystal structures: *R*^2^ = 0.34, *p* = 0.099; unknown crystal structures: *R*^2^ = 0.23, *p* = 0.019).

To conclude, good to fair and significant correlations between log(*Act*_max_) and log(*P*_EH_) of the *experimental data set* (*R*^2^ = 0.50, *p* = 4.6^*^10^−23^) and the *flexibility data set* (*R*^2^ = 0.22, *p* = 0.6^*^10^−2^) were found. Hence, promiscuous EHs have higher maximum specific activities.

### 3.7. Specific EHs prefer to hydrolyze large and flexible esters

Next, we investigated, which of the 96 esters was preferentially hydrolyzed by EHs with different *P*_EH_. As a criterion, we chose the number of freely rotatable bonds of an ester, TA **(see section 2.7)**. We did so because TA is a combined measure for an ester’s size and conformational dynamics (57). To account for the uneven distribution of esters in our data set with respect to TA, we calculated *Norm*_ester_(TA), i.e., the number of hydrolyzed esters with a specific TA (*Ester*_hydrolysed_ (TA)) divided by the total number of esters in the data set with this specific TA (*Ester*_library_(TA)) **(see section 2.7)** (Eq. 1).

According to TA, the esters were classified into 17 groups that ranged from small esters with no rotatable bond to large esters with 56 rotatable bonds (Figure 4, **Table S9)**. Esters with three (24% of the ester library) and four (16% of the ester library) rotatable bonds are most frequent. The analysis of the *experimental data set* revealed that promiscuous EHs have high *Norm*ester values irrespective of TA, i.e., promiscuous EHs accept a large variety of esters with different sizes and degrees of conformational dynamics (Figure 4A, **Table S9)**. In contrast, specific EHs only have high *Norm*ester values regarding esters with high TA, i.e., specific EHs preferentially hydrolyze (very) large and flexible esters (Figure 4A, **Table S9)**. The same tendency was observed for the *flexibility data set* (Figure 4B, **Table S9)**.

**Figure 4:**
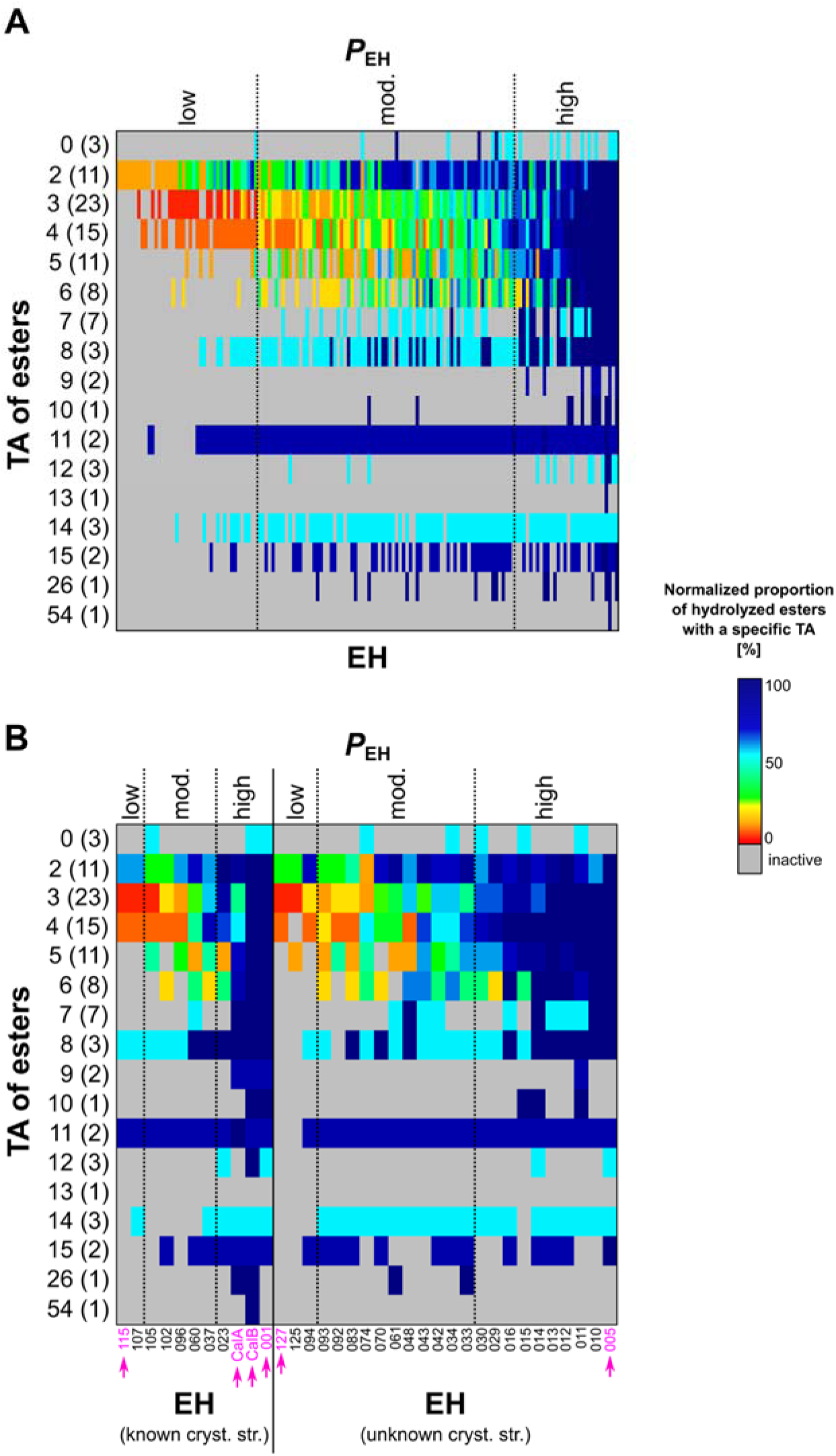
Relation between the number of esters’ TA and *P*_EH_. Relation between *Ester*norm, i.e., the relative proportion of the number of hydrolyzed esters with a specific TA, and *P*_EH_ of **(A)** the *experimental data set* and **(B)** the *flexibility data set* containing EHs with known crystal structures (left), EHs with unknown crystal structures (right), and EHs constituting the *representative data set* (indicated by magenta arrows). TA was calculated based on SMILES codes of 96 esters provided by Martínez-Martínez *et al.* (19). A blue (red) color indicates that the EH hydrolyzes many (few) esters with a specific TA relative to the total number of esters in the data set with this specific TA (see color scale on the right); the total number of esters with a specific TA is given in brackets on the y-axis. *P*_EH_ is defined as *low* if the EH hydrolyzes ≤ 9 esters, as *moderate* if the EH hydrolyzes between 10 and 29 esters, and as *high* if the EH hydrolyzes ≥ 30 esters.

To conclude, promiscuous EHs accept a large variety of esters with different sizes and degrees of conformational dynamics whereas specific EHs preferentially hydrolyze (very) large and flexible esters.

## 4. Discussion

The main outcomes of our analyses are I) that promiscuous EHs are significantly globally less flexible and have less flexible catalytically active residues than specific ones, II) that promiscuous EHs are significantly more thermostable, III) that promiscuous EHs have a significantly increased specific activity, and IV) that specific EHs prefer to hydrolyze large and flexible esters.

We established these relations using one of the still few experimental large-scale datasets where a diverse set of EHs was functionally assessed against a customized library of dissimilar esters (19). Functional promiscuity may arise from several conditions, including the environment of the enzyme or the concentration of a substrate, which may complicate the analysis of the molecular mechanism underlying promiscuity (5). Still, functional promiscuity ultimately is a result of recognition promiscuity (5); here, we therefore focused on substrate promiscuity (18), i.e., an enzyme carries out its typical catalytic function using non-canonical substrates, in that experimental conditions had been kept constant for the assessment of the different esterase/ester combinations (19). Almost all of the EHs were unambiguously assigned to one of the *F*EH of the Arpigny and Jaeger classification, which is based mainly on a comparison of amino acid sequences (53). Except for classes with a few members only (cyclase-like EHs and the yeast family), all other classes cover at least two of the three *P*_EH_ ranges such that *P*_EH_ cannot be assigned based on the EH’s class affiliation **(Figure S13, Table S10)** and, hence, amino acid sequence information. Even family *F*_IV_, which contains a higher proportion of substrate-promiscuous EHs, also contains EHs with a small substrate range.

For scrutinizing the mechanism underlying esterase promiscuity at the atomistic level, we needed to apply comparative models of EHs, since only for ~7% of the experimentally assessed EHs crystal structures were available. Restricting the generation of esterase models to sequence identities ≥ 25% with respect to available targets yielded generally good structural models both globally and locally, as also validated against cases where crystal structures are known. Throughout our study, we probed for the consistency of our analyses between subsets of EHs for which either crystal structures are known or not; we only found quantitative differences, but no qualitative ones. One of the reasons is likely that rigidity analyses were based on structural ensembles generated by multiple and μs-long MD simulations, which has been shown to improve both global and local protein structure (58, 59) to the level of approaching experimental accuracy (60) and markedly increases the robustness of the results (42). We furthermore showed that results are consistent irrespective of whether EH flexibility characteristics were assessed globally or only for CARs, and that mobility characteristics computed directly from MD trajectories show the same trend, although the correlation with *P*_EH_ is insignificant. Finally, we used experimental melting temperatures of EHs as indicators for enzyme flexibility (45, 48), which yielded the same relation with *P*_EH_ as computed flexibility characteristics. The partial use of comparative models rather than crystal structures throughout this study may lead to concern. Yet, our consistent and robust findings indicate that when applying this workflow to novel EHs, including to those for which no crystal structure exists but a structural homolog with a sequence identity ≥ 25%, it should be possible to discover enzymes with ‘sufficient’ substrate promiscuity to serve as a starting point for further exploration in biotechnology and synthetic organic chemistry. In that respect, the flexibility characteristics of EHs analyzed here have a notably stronger predictive power than the active site effective volume introduced earlier (19) (**Figure S14, Tables S11 and S12**).

The finding that promiscuous EHs are significantly globally *less* flexible and have *less* flexible catalytically active residues than specific EHs is in stark contrast to the general view of the role of structural flexibility for promiscuity (4, 5): Besides the examples of CYP and β-lactamase mentioned above, the possibility of dynamically restructuring active sites has also been recognized for other systems as underlying their promiscuity (61–64). Finally, interactions between antibodies and antigens are likely the quintessential example of the canonical relationship between flexibility and binding promiscuity: As antibodies mature to become more specific, their flexibility is decreased (5).

It has been recognized that conformational changes may not always be necessary for promiscuity if a variety of substrates can be bound by partial recognition or the presence of multiple binding sites (5). However, these cases do not seem to be relevant reasons for EH promiscuity because partial recognition often is associated with catalytic inefficiency (1), which is contrary to our observation that *P*_EH_ correlates with EH activity, and the presence of multiple binding sites that could give rise to promiscuity is controverted by the finding that promiscuous EHs have large active site effective volumes (19), i.e., large pockets with few subpockets. Inversely, our findings of rigid promiscuous EHs may be consistent with the idea that multiple ligands can be accommodated in a single site by exploiting diverse interacting residues (Figure 5).

**Figure 5:**
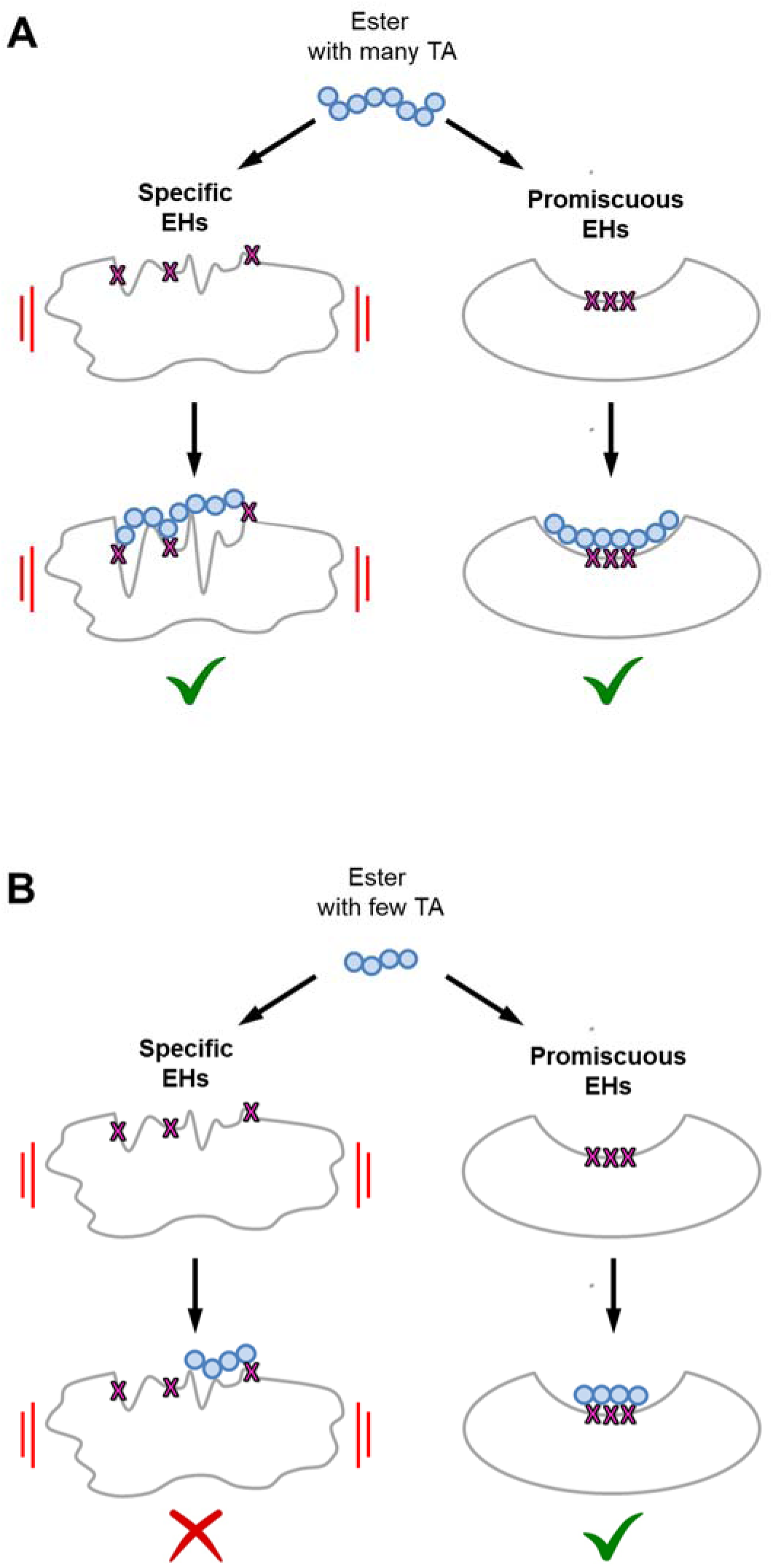
Mechanistic model of EH flexibility, ligand size and conformational dynamics affecting *P*_EH_. Impact of esters with **(A)** many or **(B)** few TA on specific, and hence more flexible (left), and promiscuous, and hence more rigid (right), EHs. Ligand parts connected by TA are represented as blue circles. Specific EHs and large ligands with many TA can mutually adapt (panel **A**, left), and promiscuous EH can bind large ligands (panel **A**, right) and small ligands (panel **B**, right) exploiting different interaction partners. Small (and/or rigid) ligands are not able to lead to a structural adaptation of specific EHs (panel **B**, left), though, resulting in conformational proofreading. The red bars indicate the flexibility of the EHs. A green tick (red cross) indicates that ester cleavage is (not) catalyzed.

Our results as to *specific but flexible* EHs may be reconciled with a model according to which conformational changes may have been selected in EH evolution for their ability to enhance specificity in recognition (Figure 5), resulting in what has been termed conformational proofreading (16). In the case of specific EHs, flexibility may help to overcome a structural mismatch between the enzyme and its substrate existing when both are in their ground states, that way enhancing recognition specificity. This view is corroborated by our finding that specific EHs prefer to hydrolyze large and flexible substrates: Larger substrates can form more interactions with the enzyme, that way helping to overcome the deformation energy required by the enzyme to optimizing the correct binding probability over the incorrect one; flexible substrates can tolerate higher strains and thus can be expected to participate in more binding events (65, 66) (Figure 5). In that respect, the relation between structural flexibility of EHs and promiscuity found here is more causative than that between active site effective volume and promiscuity (19), because small active site effective volumes found for specific EHs cannot rationalize why specific EHs prefer to hydrolyze large and flexible substrates.

In summary, the combined large-scale analysis of experimental EH promiscuity and computed EH flexibility reveals that promiscuous EHs are significantly less flexible than specific ones. This result is counterintuitive at first but may be reconciled with a model according to which multiple ligands can be accommodated in a single active site of promiscuous EHs by exploiting diverse interacting residues, whereas structural flexibility in the case of specific EHs serves for conformational proofreading. Our results furthermore signify that EH sequence space, charted, e.g., by (meta)genomics studies, can be screened by rigidity analyses for promiscuous EHs that may serve as starting points for further exploration in biotechnology and synthetic organic chemistry.

## 5. Acknowledgements

CN is funded through a grant (“Vernetzungsdoktorand”) provided by the Forschungszentrum Jülich. Parts of the study were supported by the German Federal Ministry of Education and Research (BMBF) through funding number 031B0837A “LipoBiocat” to HG and KEJ and through funding number 031L0182 “InCelluloProtStruct” to HG, the German Research Foundation (DFG) through funding no. INST 208/704-1 FUGG to HG and INST 208/654-1 FUGG to KEJ, as well as the state of North Rhine Westphalia (NRW) and the European Regional Development Fund (EFRE) through funding no. 34-EFRE-0300096 “CLIB-Kompetenzzentrum Biotechnologie (CKB)” to HG. HG is grateful for computational support and infrastructure provided by the “Zentrum für Informations-und Medientechnologie” (ZIM) at the Heinrich Heine University Düsseldorf. HG gratefully acknowledges the computing time granted by the John von Neumann Institute for Computing (NIC) and provided on the supercomputer JUWELS at Jülich Supercomputing Centre (JSC) (user IDs: HKF7; protil (project ID: 15956)) (67). HG is grateful to OpenEye for an academic license. MF acknowledges the grant ‘INMARE’ from the European Union’s Horizon 2020 (grant agreement no. 634486) and BIO2017-85522-R from the Ministerio de Ciencia, Innovación y Universidades (MCIU), Agencia Estatal de Investigación (AEI), Fondo Europeo de Desarrollo Regional (FEDER) and European Union (EU). CC thanks the Ministerio de Economía y Competitividad and FEDER for a PhD fellowship (Grant BES-2015-073829). The authors are grateful to David Almendral and Ruth Matesanz for their support of CD analysis.

## 6. Authors contributions

HG and KEJ conceived the study. CN analyzed the experimental data, performed structure prediction, MD simulations and CNA computations, analyzed the computational data, and wrote the manuscript together with HG. DM initially contributed to the structure prediction. BD performed similarity analysis of esters. MF and CC measured and analyzed melting temperatures and determined kinetic parameters. HG supervised and managed the project. All authors reviewed and approved the manuscript.

