## Supplemental Information for "Promiscuous esterases counterintuitively are less flexible than specific ones"

### Table of Contents


#### Supplemental Materials and Methods 4-8

#### Supplemental Tables 7-29

Table S1: Information about EHs of the *flexibility data set* with known crystal structures. 9

Table S2: Information about EHs of the *flexibility data set* with unknown crystal structures. 10-11

Table S3: Comparison between the *volume data set* and the *flexibility data set* regarding $P_{EH}$. 12

Table S4: Comparison between the *volume data set* and the *flexibility data set* regarding $F_{EH}$. 13

Table S5: TopScore performance on comparative models of EHs of the *flexibility data set* with known crystal structures. 14

Table S6: $RMSF_{EH}$ and $RMSF_{CAR}$ of the *representative data set*. 15

Table S7: Melting temperatures of EHs determined by CD spectroscopy. 16

Table S8: $P_{EH}$, log($P_{EH}$), $Act_{max}$, and log($Act_{max}$) of EHs. 17-20

Table S9: Ester library classified according to TA. 21-23

Table S10: Distribution of $P_{EH}$ in $F_{EH}$ of the *experimental data set*. 24-27

Table S11: $P_{EH}$ and $Vol_{eff}$ of comparative models of EHs of the *flexibility data set* with known crystal structures. 28

Table S12: $P_{EH}$ and $Vol_{eff}$ of comparative models of EHs of the *flexibility data set* without known crystal structures. 29

#### Supplemental Figures 30-44

Figure S1: Correlation of the promiscuity index *I* versus $P_{EH}$. 30

Figure S2: Correlation of the number of substrates with $k_{cat}$ / $K_m$ > 0.05 $min^{-1}$ $M^{-1}$ versus $P_{EH}$. 31

Figure S3: Hierarchical clustering of ester substrates. 32

Figure S4: Distribution of the mean Tanimoto-Combo similarity scores $\delta_i$. 33

Figure S5: Comparison between the *volume data set* and the *flexibility data set* regarding $P_{EH}$. 34

Figure S6: Comparison between the *volume data set* and the *flexibility data set* regarding $F_{EH}$. 35

Figure S7: TopScore performance on comparative models of EHs of the *flexibility data set* with known crystal structures. 36

Figure S8: Substrate-accessibility of EHs of the *representative data set*. 37

Figure S9: Correlation of $RMSF_{EH}$ or $T_p$ *versus* $P_{EH}$ of the *representative data set*. 38

Figure S10: Correlation of $Flex_{CAR}$ *versus* $P_{EH}$. 39-40

Figure S11: Correlation of $RMSF_{CAR}$ or $Flex_{CAR}$ *versus* $P_{EH}$ of the *representative data set*. 41

Figure S12: Correlation of log($Act_{max}$) *versus* log($P_{EH}$). 42

Figure S13: Distribution of $P_{EH}$ in $F_{EH}$ of the *experimental data set*. 43

Figure S14: Correlation of $Vol_{eff}$ *versus* $P_{EH}$. 44

**Supplemental References** 45-47

### Supplemental Materials and Methods

##### Determining and classifying $P_{EH}$

Martínez-Martínez *et al.* (1) examined $P_{EH}$ of all esterases (EHs) with a kinetic pH indicator assay (2-4), which unambiguously allows quantifying specific activities (*Act*) at pH 8.0 and 30 °C, using a substrate concentration above 0.5 mM. The specific activities were given in units (U) / (g wet cells); for CalA and CalB preparations, the specific activities were given in U / (g total protein). The assays were performed as triplicates, with the average specific activity ($Act_{average}$) given and standard deviation (STD) ≤ 1% in all cases. Additionally, the average maximum specific activity ($Act_{max}$) was determined for each EH.

In order to rank (classify) $P_{EH}$, the authors introduced a structural parameter, the active site effective volume ($Vol_{eff}$). $Vol_{eff}$ represents the topology of the active site in terms of the active site cavity volume ($Vol_{cav}$) computed by Fpocket (5) divided by the relative solvent-accessible surface area ($SASA_{rel}$) using GetArea Web server (6) **(Eq. S1)**.

$$Vol_{eff}[\text{Å}^3] = \frac{Vol_{cav}\,[\text{Å}^3]}{SASA_{rel}} \qquad \textbf{Eq. S1}$$

$Vol_{eff}$ was computed for 96 EHs (termed *volume data set*) for which the following four criteria were satisfied:

I. Eleven EHs with known crystal structures were included.

II. Homology models of 85 EHs with unknown crystal structures were generated using the Prime software from Schrödinger (7) (known crystal structures from EHs in I were used as templates).

III. EHs in II showed sequence identities ≥ 25% (in comparison to known crystal structures from EHs in I).

IV. Catalytically active residues (CARs) were unambiguously identified.

##### Promiscuity index

Functional promiscuity ultimately is a result of recognition promiscuity (8). Here, we therefore focused on substrate promiscuity (9), i.e., an enzyme carries out its typical catalytic function using non-canonical substrates, in that experimental conditions had been kept constant for the assessment of the different esterase/ester combinations (1). Under these conditions, an entropy-based metric can be adapted to quantify substrate promiscuity (10). With the conventional definition of the catalytic efficiency $e$ (**Eq. S2**) (11)

$$e = k_{cat} / K_m \qquad \textbf{Eq. S2,}$$

which ideally serves as the kinetic parameter in enzyme promiscuity studies for comparison (10, 12), $p_i$ can be conceptualized as the probability that the $i^{th}$ substrate will be the first to be metabolized when an enzyme is simultaneously exposed to equal, low concentrations of all $N$ substrates (**Eq. S3**) (10).

$$p_i = e_i / \sum_{i=1}^{N} e_i \qquad \textbf{Eq. S3}$$

Then, applying the concept of information entropy, a promiscuity index $I$ can be defined (**Eq. S4**) (10):

$$I = -\frac{1}{\log N} \sum_{i=1}^{N} p_i \log p_i \qquad \textbf{Eq. S4}$$

The lower limit of $I$ is 0 if the enzyme only turns over one substrate, i.e., is perfectly specific; it is 1 if the enzyme equally-well metabolizes all substrates, i.e., is perfectly promiscuous.

**Assessing similarities of ester substrates**

For assessing the similarity of the ester substrates, the maximum pairwise Tanimoto-Combo similarity scores $\delta_{ij}$ for compound $i$ versus $j$ were calculated using ROCS (13) (ROCS 3.3.2.2: OpenEye Scientific Software, Santa Fe, NM.). These scores range from 0, for dissimilar molecules, to 2, for identical molecules. Prior to their calculation, an energy-minimized 3D structure of each of the 96 compounds of interest (queries) was generated using LigPrep (Schrödinger Release 2020-2: Schrödinger, LLC, New York, NY, 2020). This structure was aligned then against a conformational database of 3D conformers of all 96 compounds. For each pair of compounds, the maximum Tanimoto-Combo score is attributed to the best pairwise alignment between a given compound query and the most similar conformer of another compound in the conformational database. The 3D conformational database was prepared using OMEGA (14) (OMEGA 3.0.0.1: OpenEye Scientific Software, Santa Fe, NM.) with at most 200 conformers generated per compound. Finally, the mean maximum pairwise Tanimoto-Combo similarity score $\delta_i$ of a substrate $i$ to all other substrates in the data set was computed.

**Model quality assessment by TopScore and validation**

TopScore (15) predicts 1 – lDDT, with lDDT being the local Distance Difference Test (16), a measure for structural similarity that does not require superimpositioning of two structures. Therefore, the range of TopScore is [0, 1], with 0 (1) indicating low (high) estimated errors of the residues and models.

For validation, 1 - lDDT was also computed for EHs with known crystal structure and the respective comparative model, using the lDDT web server from Swiss-Model (16). Note that in this case, the comparative model was generated by TopModel excluding the known crystal structure.

**Starting structure preparation, parametrization, and equilibration**

The EH structures were preprocessed with the Protein Preparation Wizard of Schrödinger's Maestro Suite (17). Of a crystal structure, we used only chain A and removed structurally resolved water molecules and ligands. Non-resolved termini were connected to acetyl (ACE) and *N*-methyl amide (NME) groups to avoid artificially charged termini. In order to match the experimental conditions of pH 8.0 **(see section 2.1)**, we used Epik (18) to calculate the p*K*a of relevant functional groups. All hydrogen atoms were then added according to the Amber ff14SB library (19). The prepared EH structures were solvated with OPC water (20), leaving at least 12 Å between the EH structure and the edges of the solvent box, by using LeaP of Amber19 (21). We also added sodium counter ions to ensure the neutrality of the system.

We used the Amber ff14SB force field (19) to parametrize the protein. Ion parameters were taken from Joung and Cheatham (22). The detailed minimization, thermalization, and equilibration protocol has been reported in ref. (23). In short, the system was initially subjected to three rounds of energy minimization to get rid of any bad contacts. The system was heated to 300 K and the pressure was adapted such that a density of 1 g $cm^{-3}$ was obtained. During thermalization and density adaptation, we kept the solute fixed by positional restraints of 1 kcal $mol^{-1}$ $Å^{-2}$, which were gradually removed. Subsequently, the system was subjected to five independent NPT production simulations of 1 µs length each using unbiased MD simulations. Therefore, the initial velocities were randomly assigned during the first step of the production simulations.

**Thermal unfolding simulations**

Therefore, a hydrogen bond energy $E_{HB}$ is computed by a modified version of the potential by Mayo *et al.* (24). For a given network state $\sigma = f(T)$, hydrogen bonds with an energy $E_{HB}$ >

$E_{cut}(\sigma)$ are removed from the network at temperature $T$. In the present study, thermal unfolding simulations were carried out by decreasing $E_{cut}$ from -0.1 kcal mol$^{-1}$ to -6.0 kcal mol$^{-1}$ with a step size of 0.1 kcal mol$^{-1}$. As $E_{cut}$ can be converted to a temperature $T$ using the linear equation introduced by Radestock *et al.* (25, 26) **(Eq. S5)**, the range of $E_{cut}$ is equivalent to increasing the temperature from 302 K to 420 K with a step size of 2 K. Along the thermal unfolding simulations, hydrophobic interactions were not removed because they remain constant in strength or become even stronger with increasing temperature (27).

$$T = \frac{-20\ \mathrm{K}}{\mathrm{kcal} \cdot \mathrm{mol}^{-1}} E_{\mathrm{cut}} + 300\ \mathrm{K} \qquad \textbf{Eq. S5}$$

**Cluster configuration entropy and stability map**

The cluster configuration entropy $H_{type2}$ was used to identify the phase transition temperature $T_p$ of the EHs constituting the *flexibility data set* during the thermal unfolding simulation. At $T_p$, the protein switches from a rigid (structurally stable) to a floppy (unfolded) state. However, the percolation behavior of protein networks is usually more complex, and multiple phase transitions can be observed (25, 26, 28-32). Initially, the protein network is dominated by a giant rigid cluster, and $H_{type2}$ is low because of the limited number of possible ways to configure a system with this cluster. When the giant rigid cluster starts to decay or stops to dominate the network, $H_{type2}$ jumps. There, the network is in a partially flexible state with many ways to configure a system consisting of many small clusters. In order to determine $T_p$, a double sigmoid fit was applied to an $H_{type2}$ *versus* $T(E_{cut})$ curve as done previously (25, 26, 28-32), and $T_p$ taken as that $T$ value associated with the largest slope of the fit. The rigid cluster decomposition of the EHs was visually inspected by VisualCNA (33), an easy-to-use PyMOL plug-in that allows setting up CNA runs and analyzing CNA results linking data plots with molecular graphics representations. VisualCNA is available under an academic license from https://cpclab.uni-duesseldorf.de/index.php/Software.

During a thermal unfolding simulation, the stability map $rc_{ij}$ indicates for all residue pairs the $E_{cut}$ value at which a rigid contact $rc$ between the two residues $i$ and $j$ (represented by their $C_\alpha$ atoms) is lost; $rc$ exists as long as $i$ and $j$ belong to the same rigid cluster $c$ of the set of rigid clusters $C^{E_{cut}}$ (34). Thus, $rc_{ij}$ contains information about the rigid cluster decomposition cumulated over all network states $\sigma$ during the thermal unfolding simulation. The sum over all entries in $rc_{ij}$ yields the chemical potential energy due to non-covalent bonding, obtained from the coarse-grained, residue-wise network representation of the underlying protein structure

(29). In the present study, we applied the neighbor stability map $rc_{ij,neighbor}$ of each EH to investigate short-range rigid contacts. For this, as done previously (29, 32), $rc_{ij}$ was filtered such that only rigid contacts between two residues that are at most 5 Å apart from each other were considered. Here, in particular, we focused on rigid contacts between CARs and other residues at most 5 Å apart and calculated the average over all such entries in $rc_{ij,neighbor}$ (termed $Flex_{\mathrm{CAR}}$).

### Supplemental Tables

##### Table S1: Information about EHs of the *flexibility data set* with known crystal structures.

| EH | PDB | $P_{EH}$[a] | $F_{EH}$[b] | Global Topscore[c] | CARs[d] | $T_p$ [K][e] | $Flex_{CAR}$ [kcal/mol][f] |
|---|---|---|---|---|---|---|---|
| 001 | 5JD4_A | 72 | IV | 0.0849 | S161; D256; H286 | 357.19 ± 0.46 | -1.91 ± 0.05 |
| CalB | 4K6G_A | 68 | Yeast | 0.0744 | S107; D189; H226 | 351.60 ± 0.51 | -1.95 ± 0.07 |
| CalA147 | 3GUU_A | 36 | Yeast | 0.1739 | S205; D355; H387 | 346.17 ± 1.36 | -1.31 ± 0.06 |
| 023 | 4Q3O_A | 34 | IV | 0.1855 | S194; D290; H320 | 329.31 ± 0.89 | -1.23 ± 0.02 |
| 037 | 5JD5_A | 28 | IV | 0.1318 | S169; D265; H295 | 345.27 ± 1.29 | -1.15 ± 0.05 |
| 060 | 4I3F_A | 21 | C-C MCPh | 0.0968 | S104; D230; H258 | 345.82 ± 0.87 | -1.41 ± 0.04 |
| 096 | 4FBM_A | 11 | V | 0.2114 | S126; D227; H257 | 344.31 ± 0.57 | -1.06 ± 0.06 |
| 102 | 5JD3_A | 10 | II | 0.1010 | S15; D192; H195 | 333.18 ± 0.93 | -1.46 ± 0.07 |
| 105 | 5IBZ_A | 10 | Cyclase-like | 0.3046 | F84; R87; Q127; Q131; D133; H137; H286; E299 | 341.19 ± 0.91 | -0.66 ± 0.03 |
| 107 | 4Q3L_A | 9 | V | 0.1181 | S97; D221; H249 | 336.34 ± 0.35 | -1.36 ± 0.04 |
| 115 | 4Q3K_A | 8 | I | 0.1436 | S113; D169; H201 | 322.39 ± 0.65 | -0.74 ± 0.05 |

[a] Experimentally measured substrate promiscuity level of EHs provided by Martínez-Martínez *et al.* (1) **(see section 2.1)**.

[b] EH families based on the Arpigny and Jaeger classification (35) **(see section 2.1)**.

[c] Whole-protein error estimates predicted by TopScore for comparative models (15) **(see section 2.2)**.

[d] Catalytically active residues of EHs.

[e] Global flexibilities of EHs with SEM based on predicted $H_{type2}$ **(see section 2.5)**.

[f] Local flexibilities of catalytically active residues of EHs with SEM based on predicted $rc_{ij,neighbor}$ **(see section 2.5)**.

**Table S2: Information about EHs of the *flexibility data set* with unknown crystal structures.**

| EH | $P_{EH}$[a] | $F_{EH}$[b] | Global TopScore[c] | CARs[d] | $T_p$ [K][e] | $Flex_{CAR}$ [kcal/mol][f] |
|---|---|---|---|---|---|---|
| 005 | 67 | IV | 0.0945 | S159; D254; H284 | 351.10 ± 0.88 | -1.86 ± 0.05 |
| 010 | 58 | IV | 0.1586 | S181; D279; H309 | 358.55 ± 0.38 | -1.50 ± 0.05 |
| 011 | 53 | IV | 0.0878 | S159; D254; H284 | 349.80 ± 1.25 | -1.34 ± 0.05 |
| 012 | 51 | IV | 0.0939 | S159; D254; H284 | 351.98 ± 1.35 | -1.66 ± 0.06 |
| 013 | 49 | IV | 0.1325 | S171; D268; H298 | 348.61 ± 0.45 | -1.24 ± 0.06 |
| 014 | 48 | IV | 0.1840 | S190; D290; H320 | 348.72 ± 0.88 | -1.42 ± 0.05 |
| 015 | 42 | IV | 0.1123 | S146; E240; H270 | 338.14 ± 0.84 | -1.40 ± 0.06 |
| 016 | 42 | IV | 0.0934 | S159; D254; H284 | 335.59 ± 1.87 | -1.87 ± 0.05 |
| 029 | 31 | IV | 0.0869 | S159; D254; H284 | 328.91 ± 0.89 | -1.48 ± 0.06 |
| 030 | 30 | VI | 0.1118 | S116; D164 ; H195 | 342.22 ± 0.75 | -0.67 ± 0.06 |
| 033 | 29 | C-C MCPh | 0.1514 | S104; D225; H253 | 344.20 ± 0.84 | -1.26 ± 0.06 |
| 034 | 29 | VI | 0.1505 | S119; D173; H204 | 341.87 ± 1.87 | -1.01 ± 0.06 |
| 042 | 27 | IV | 0.1065 | S125; E218; H248 | 341.16 ± 1.12 | -1.21 ± 0.04 |
| 043 | 27 | IV | 0.1050 | S144; E238; H268 | 338.57 ± 1.24 | -0.72 ± 0.04 |
| 048 | 23 | IV | 0.1020 | S146; E240; H270 | 340.73 ± 1.14 | -1.13 ± 0.05 |
| 061 | 20 | VI | 0.1414 | S118; D172; H203 | 339.97 ± 0.93 | -0.66 ± 0.05 |
| 070 | 18 | CE | 0.1273 | S189; D279; H308 | 328.37 ± 0.36 | -1.09 ± 0.05 |
| 074 | 17 | V | 0.1056 | S94; D203; H231 | 332.59 ± 1.12 | -1.05 ± 0.05 |

| | | | | | | |
|---|---|---|---|---|---|---|
| 083 | 14 | V | 0.2488 | S120; D247; H275 | 335.31 ± 3.01 | -0.68 ± 0.05 |
| 092 | 12 | V | 0.2161 | S105; D233; H261 | 324.94 ± 1.44 | -1.12 ± 0.05 |
| 093 | 12 | VII | 0.2687 | S183; D311; H420 | 320.43 ± 1.13 | -1.30 ± 0.06 |
| 094 | 12 | V | 0.2396 | S127; D246; H279 | 330.70 ± 1.74 | -0.70 ± 0.05 |
| 125 | 4 | / | 0.1248 | S70; D149; H174 | 328.66 ± 1.73 | -0.94 ± 0.03 |
| 127 | 4 | V | 0.2367 | S101; D236; H263 | 318.60 ± 1.57 | -1.10 ± 0.05 |

[a] Experimentally measured substrate promiscuity level of EHs provided by Martínez-Martínez *et al.* (1) **(see section 2.1)**.

[b] EH families based on the Arpigny and Jaeger classification (35) **(see section 2.1)**.

[c] Whole-protein error estimates predicted by TopScore for the comparative models (15) **(see section 2.2)**.

[d] Catalytically active residues of EHs.

[e] Global flexibilities of EHs with SEM based on predicted $H_{\text{type2}}$ **(see section 2.5)**.

[f] Local flexibilities of catalytically active residues of EHs with SEM based on predicted $rc_{ij,neighbor}$ **(see section 2.5)**.

**Table S3: Comparison between the *volume data set* and the *flexibility data set* regarding $P_{EH}$.**

| $P_{EH}$[a] | #EHs of the *volume data set*[b] | #EHs of the *flexibility data set*[c] |
|---|---|---|
| Low | 19 (19.79) | 4 (11.43) |
| Moderate | 51 (53.13) | 17 (48.57) |
| High | 26 (27.08) | 14 (40.00) |
| **#EHs** | **96** | **35** |

[a] Experimentally measured substrate promiscuity level of EHs provided by Martínez-Martínez *et al.* (1) **(see section 2.1)**. $P_{EH}$ is defined as *low* if the EH hydrolyzes ≤ 9 esters, as *moderate* if the EH hydrolyzes between 10 and 29 esters, and as *high* if the EH hydrolyzes ≥ 30 esters.

[b] Values in brackets represent the relative proportions of EHs in the *volume data set* in %.

[c] Values in brackets represent the relative proportions of EHs in the *flexibility data set* in %.

**Table S4: Comparison between the *volume data set* and the *flexibility data set* regarding $F_{EH}$.**

| $F_{EH}$[a] | #EHs of the *volume data set*[b] | #EHs of the *flexibility data set*[c] |
|---|---|---|
| FI | 6 (6.52) | 1 (2.94) |
| FII | 7 (7.61) | 1 (2.94) |
| FIV | 32 (34.78) | 15 (44.12) |
| FV | 23 (25.00) | 7 (20.59) |
| FVI | 5 (5.43) | 3 (8.82) |
| FVII | 4 (4.35) | 1 (2.94) |
| CE | 3 (3.26) | 1 (2.94) |
| C-C MCPh | 9 (9.78) | 2 (5.88) |
| Cyclase-like | 1 (1.09) | 1 (2.94) |
| Yeast class | 2 (2.17) | 2 (5.88) |
| Unclassified | 4 (4.35) | 1 (2.94) |
| **#EHs** | **96** | **35** |

[a] EH families based on the Arpigny and Jaeger classification (35) **(see section 2.1)**.

[b] Values in brackets represent the relative proportions of EHs in the *volume data set* in %.

[c] Values in brackets represent the relative proportions of EHs in the *flexibility data set* in %.

**Table S5: TopScore performance on comparative models of EHs of the *flexibility data set* with known crystal structures.**

| EH | PDB ID[a] | Global TopScore[b] | 1 - lDDT score[c] |
|---|---|---|---|
| 001 | 5JD4_A | 0.1501 | 0.2019 |
| CalB | 4K6G_A | 0.0958 | 0.0998 |
| CalA | 3GUU_A | 0.2219 | 0.1139 |
| 023 | 4Q3O_A | 0.2578 | 0.1976 |
| 037 | 5JD5_A | 0.2291 | 0.2622 |
| 060 | 4I3F_A | 0.1644 | 0.195 |
| 096 | 4FBM_A | 0.1997 | 0.146 |
| 102 | 5JD3_A | 0.0954 | 0.1376 |
| 105 | 5IBZ_A | 0.3519 | 0.6241 |
| 107 | 4Q3L_A | 0.1798 | 0.2249 |
| 115 | 4Q3K_A | 0.1788 | 0.2128 |

[a] PDB IDs that were used as references to calculate the 1 – lDDT (local Distance Difference Test) scores.

[b] Whole-protein error estimates predicted by TopScore (15) **(see section 2.2)**.

[c] Local Distance Difference Test computed by the Swiss-Model web server (16) **(see section 2.2)**.

**Table S6: *RMSF*$_{EH}$ and *RMSF*$_{CAR}$ of the *representative data set*.**

| EH | *RMSF*$_{EH}$ [Å][a] | *RMSF*$_{CAR}$ [Å][b] |
|---|---|---|
| 115 | 1.27 ± 0.03 | 1.60 ± 0.22 |
| 001 | 0.91 ±0.01 | 0.60 ± 0.03 |
| 127 | 1.54 ± 0.04 | 1.03 ± 0.06 |
| 005 | 1.13 ± 0.02 | 0.70 ± 0.04 |
| CalA | 1.76 ± 0.04 | 1.03 ± 0.07 |
| CalB | 0.90 ± 0.02 | 0.62 ± 0.02 |

[a] Average per-residue root-mean-square fluctuations for EHs **(see section 2.6)**.

[b] Average per-residue root-mean-square fluctuations for catalytically active residues of EHs **(see section 2.6)**.

**Table S7: Melting temperatures of EHs determined by CD spectroscopy.**

| EH | $P_{EH}$[a] | $T_d$ [°C][b] |
|---|---|---|
| 001 | 72 | 42.10 ± 0.20 |
| 002 | 71 | 47.45 ± 0.31 |
| 003 | 69 | 45.90 ± 0.43 |
| 004 | 67 | 44.63 ± 0.19 |
| 006 | 66 | 58.57 ± 0.24 |
| 008 | 63 | 38.31 ± 0.44 |
| 009 | 61 | 36.10 ± 0.73 |
| 016 | 42 | 35.92 ± 0.69 |
| 021 | 36 | 35.31 ± 0.62 |
| 037 | 28 | 35.99 ± 0.20 |
| 043 | 27 | 39.98 ± 0.74 |

[a] Experimentally measured substrate promiscuity level of EHs provided by Martínez-Martínez *et al.* (1) **(see section 2.1)**.

[b] Melting temperatures of EHs ± STD ($n$ = 3) determined by CD spectroscopy **(see section 2.8)**.

**Table S8: $P_{EH}$, log($P_{EH}$), $Act_{max}$, and log($Act_{max}$) of EHs.**

| EH[a] | $P_{EH}$[b] | log($P_{EH}$) | $Act_{max}$ [U / (g wet cells)][c] | log($Act_{max}$) [log(U / (g wet cells))] |
|---|---|---|---|---|
| **001** | 72 | 1.86 | 1326.63 | 3.12 |
| 002 | 71 | 1.85 | 113.26 | 2.05 |
| 003 | 69 | 1.84 | 106.35 | 2.03 |
| **CalB** | 68 | 1.83 | 69105.06 [U / (g total protein)] | n.d.[d] |
| 004 | 67 | 1.83 | 262.23 | 2.42 |
| **005** | 67 | 1.83 | 23.52 | 1.37 |
| 006 | 66 | 1.82 | 338.55 | 2.53 |
| 007 | 64 | 1.81 | 77.72 | 1.89 |
| 008 | 63 | 1.80 | 2239.16 | 3.35 |
| 009 | 61 | 1.79 | 168.40 | 2.23 |
| **010** | 58 | 1.76 | 77.55 | 1.89 |
| **011** | 53 | 1.72 | 120.02 | 2.08 |
| **012** | 51 | 1.71 | 137.81 | 2.14 |
| **013** | 49 | 1.69 | 278.08 | 2.44 |
| **014** | 48 | 1.68 | 138.13 | 2.14 |
| **015** | 42 | 1.62 | 93.93 | 1.97 |
| **016** | 42 | 1.62 | 991.93 | 3.00 |
| 017 | 39 | 1.59 | 7787.23 | 3.89 |
| 018 | 38 | 1.58 | 304.25 | 2.48 |
| 019 | 37 | 1.57 | 5038.96 | 3.70 |
| 020 | 37 | 1.57 | 35.12 | 1.55 |
| 021 | 36 | 1.56 | 963.46 | 2.98 |
| **CalA** | 36 | 1.56 | 25224.17 [U / (g total protein)] | n.d.[d] |
| 022 | 35 | 1.54 | 1366.25 | 3.14 |
| **023** | 34 | 1.53 | 6005.66 | 3.78 |
| 024 | 34 | 1.53 | 123.42 | 2.09 |
| 025 | 33 | 1.52 | 1441.93 | 3.16 |
| 026 | 32 | 1.51 | 50.93 | 1.71 |
| 027 | 32 | 1.51 | 90.19 | 1.96 |
| 028 | 31 | 1.49 | 667.07 | 2.82 |
| **029** | 31 | 1.49 | 7660.87 | 3.88 |
| **030** | 30 | 1.48 | 752.11 | 2.88 |
| 031 | 29 | 1.46 | 398.62 | 2.60 |
| 032 | 29 | 1.46 | 242.56 | 2.38 |
| **033** | 29 | 1.46 | 376.69 | 2.58 |
| **034** | 29 | 1.46 | 1207.80 | 3.08 |
| 035 | 29 | 1.46 | 32.65 | 1.51 |
| 036 | 28 | 1.45 | 311.41 | 2.49 |
| **037** | 28 | 1.45 | 746.72 | 2.87 |
| 038 | 28 | 1.45 | 193.26 | 2.29 |
| 039 | 28 | 1.45 | 39.50 | 1.60 |
| 040 | 27 | 1.43 | 81.25 | 1.91 |

| | | | | |
|---|---|---|---|---|
| 041 | 27 | 1.43 | 1198.35 | 3.08 |
| **<u>042</u>** | 27 | 1.43 | 139.81 | 2.15 |
| **<u>043</u>** | 27 | 1.43 | 571.48 | 2.76 |
| 044 | 25 | 1.40 | 101.91 | 2.01 |
| 045 | 24 | 1.38 | 143.52 | 2.16 |
| 046 | 23 | 1.36 | 20.79 | 1.32 |
| 047 | 23 | 1.36 | 274.96 | 2.44 |
| **<u>048</u>** | 23 | 1.36 | 148.60 | 2.17 |
| 049 | 23 | 1.36 | 9.92 | 1.00 |
| 050 | 22 | 1.34 | 661.50 | 2.82 |
| 051 | 22 | 1.34 | 278.89 | 2.45 |
| 052 | 21 | 1.32 | 252.19 | 2.40 |
| 053 | 21 | 1.32 | 90.48 | 1.96 |
| 054 | 21 | 1.32 | 2665.38 | 3.43 |
| 055 | 21 | 1.32 | 20.54 | 1.31 |
| 056 | 21 | 1.32 | 19.52 | 1.29 |
| 057 | 21 | 1.32 | 440.22 | 2.64 |
| 058 | 21 | 1.32 | 348.91 | 2.54 |
| 059 | 21 | 1.32 | 17.96 | 1.25 |
| **060** | 21 | 1.32 | 240.62 | 2.38 |
| **<u>061</u>** | 20 | 1.30 | 621.61 | 2.79 |
| 062 | 20 | 1.30 | 243.06 | 2.39 |
| 063 | 20 | 1.30 | 197.90 | 2.30 |
| 064 | 20 | 1.30 | 34.38 | 1.54 |
| 065 | 20 | 1.30 | 18.26 | 1.26 |
| 066 | 19 | 1.28 | 101.69 | 2.01 |
| 067 | 18 | 1.26 | 124.41 | 2.09 |
| 068 | 18 | 1.26 | 49.73 | 1.70 |
| 069 | 18 | 1.26 | 114.76 | 2.06 |
| **<u>070</u>** | 18 | 1.26 | 22.16 | 1.35 |
| 071 | 18 | 1.26 | 89.21 | 1.95 |
| 072 | 18 | 1.26 | 189.15 | 2.28 |
| 073 | 17 | 1.23 | 677.95 | 2.83 |
| **<u>074</u>** | 17 | 1.23 | 25.25 | 1.40 |
| 075 | 16 | 1.20 | 93.59 | 1.97 |
| 076 | 16 | 1.20 | 131.29 | 2.12 |
| 077 | 16 | 1.20 | 349.92 | 2.54 |
| 078 | 15 | 1.18 | 195.57 | 2.29 |
| 079 | 14 | 1.15 | 16.12 | 1.21 |
| 080 | 14 | 1.15 | 120.56 | 2.08 |
| 081 | 14 | 1.15 | 40.62 | 1.61 |
| 082 | 14 | 1.15 | 8978.87 | 3.95 |
| **<u>083</u>** | 14 | 1.15 | 155.38 | 2.19 |
| 084 | 13 | 1.11 | 273.06 | 2.44 |
| 085 | 13 | 1.11 | 69.15 | 1.84 |
| 086 | 13 | 1.11 | 25.46 | 1.41 |
| 087 | 13 | 1.11 | 11.45 | 1.06 |
| 088 | 13 | 1.11 | 4646.55 | 3.67 |
| 089 | 13 | 1.11 | 62.73 | 1.80 |

| | | | | |
|---|---|---|---|---|
| 090 | 13 | 1.11 | 15.24 | 1.18 |
| 091 | 13 | 1.11 | 243.86 | 2.39 |
| **<u>092</u>** | 12 | 1.08 | 41.08 | 1.61 |
| **<u>093</u>** | 12 | 1.08 | 466.48 | 2.67 |
| **<u>094</u>** | 12 | 1.08 | 41.73 | 1.62 |
| 095 | 11 | 1.04 | 98.26 | 1.99 |
| **096** | 11 | 1.04 | 5.65 | 0.75 |
| 097 | 11 | 1.04 | 191.44 | 2.28 |
| 098 | 11 | 1.04 | 24.79 | 1.39 |
| 099 | 11 | 1.04 | 498.13 | 2.70 |
| 100 | 11 | 1.04 | 241.38 | 2.38 |
| 101 | 11 | 1.04 | 17.89 | 1.25 |
| **102** | 10 | 1.00 | 3328.23 | 3.52 |
| 103 | 10 | 1.00 | 91.17 | 1.96 |
| 104 | 10 | 1.00 | 56.63 | 1.75 |
| **105** | 10 | 1.00 | 45.59 | 1.66 |
| 106 | 9 | 0.95 | 11.56 | 1.06 |
| **107** | 9 | 0.95 | 16.33 | 1.21 |
| 108 | 9 | 0.95 | 159.45 | 2.20 |
| 109 | 9 | 0.95 | 17.65 | 1.25 |
| 110 | 8 | 0.90 | 332.72 | 2.52 |
| 111 | 8 | 0.90 | 13.97 | 1.15 |
| 112 | 8 | 0.90 | 312.09 | 2.49 |
| 113 | 8 | 0.90 | 11.00 | 1.04 |
| 114 | 8 | 0.90 | 13.15 | 1.12 |
| **115** | 8 | 0.90 | 148.37 | 2.17 |
| 116 | 7 | 0.85 | 19.84 | 1.30 |
| 117 | 6 | 0.78 | 4.22 | 0.63 |
| 118 | 6 | 0.78 | 29.01 | 1.46 |
| 119 | 6 | 0.78 | 25.83 | 1.41 |
| 120 | 5 | 0.70 | 9.15 | 0.96 |
| 121 | 5 | 0.70 | 131.87 | 2.12 |
| 122 | 5 | 0.70 | 3.35 | 0.53 |
| 123 | 4 | 0.60 | 8.15 | 0.91 |
| 124 | 4 | 0.60 | 21.63 | 1.34 |
| **<u>125</u>** | 4 | 0.60 | 6.31 | 0.80 |
| 126 | 4 | 0.60 | 4.65 | 0.67 |
| **<u>127</u>** | 4 | 0.60 | 4.59 | 0.66 |
| 128 | 4 | 0.60 | 11.63 | 1.07 |
| 129 | 3 | 0.48 | 7.32 | 0.86 |
| 130 | 2 | 0.30 | 23.86 | 1.38 |
| 131 | 2 | 0.30 | 4.16 | 0.62 |
| 132 | 2 | 0.30 | 4.86 | 0.69 |
| 133 | 2 | 0.30 | 3.67 | 0.56 |
| 134 | 2 | 0.30 | 1.73 | 0.24 |
| 135 | 2 | 0.30 | 3.94 | 0.60 |
| 136 | 2 | 0.30 | 3.29 | 0.52 |
| 137 | 2 | 0.30 | 3.32 | 0.52 |
| 138 | 2 | 0.30 | 2.83 | 0.45 |

| | | | | |
|---|---|---|---|---|
| 139 | 2 | 0.30 | 4.16 | 0.62 |
| 140 | 1 | 0.00 | 1.73 | 0.24 |
| 141 | 1 | 0.00 | 2.55 | 0.41 |
| 142 | 1 | 0.00 | 0.25 | -0.59 |
| 143 | 1 | 0.00 | 1.31 | 0.12 |
| 144 | 1 | 0.00 | 1.80 | 0.25 |
| 145 | 1 | 0.00 | 2.48 | 0.39 |

[a] EHs highlighted in bold constitute the *flexibility data set*; for underlined EHs, no crystal structure is known.

[b] Experimentally determined substrate promiscuity level of EHs provided by Martínez-Martínez *et al.* (1) **(see section 2.1)**.

[c] Experimentally determined average maximum specific activities of EHs provided by Martínez-Martínez *et al.* (1) **(see section 2.1)**.

[d] Not determined.

**Table S9: Ester library classified according to TA.**

| Ester | TA |
|---|---|
| ϒ-Valerolactone | 0 |
| D-Pantolactone | 0 |
| L-Pantolactone | 0 |
| 1-Napthyl acetate | 2 |
| Ethyl acetate | 2 |
| Methyl 3-hydroxybenzoate | 2 |
| Methyl 2-hydroxybenzoate | 2 |
| Methyl benzoate | 2 |
| Vinyl acetate | 2 |
| Methyl glycolate | 2 |
| (+)-Methyl D-Lactate | 2 |
| (-)-Methyl L-Lactate | 2 |
| Phenyl acetate | 2 |
| Glyceryl trilaurate | 2 |
| Ethyl propionate | 3 |
| Ethyl benzoate | 3 |
| (1*R*)-(-)-Menthyl acetate | 3 |
| (1*S*)-(+)-Menthyl acetate | 3 |
| Methyl (*R*)-(-)-mandelate | 3 |
| Methyl (*S*)-(+)-mandelate | 3 |
| (+)-Ethyl D-Lactate | 3 |
| (−)-Ethyl L-lactate | 3 |
| (+)-Methyl (*S*)-3-hydroxybutyrate | 3 |
| (-)-Methyl (*R*)-3-hydroxybutyrate | 3 |
| (1R)-(+)-Neomenthyl acetate | 3 |
| (1S)-(+)-Neomenthyl acetate | 3 |
| Methyl butyrate | 3 |
| Methyl 2,5-dihydroxycinnamate | 3 |
| Methyl cinnamate | 3 |
| Vinyl propionate | 3 |
| Vinyl benzoate | 3 |
| Vinyl crotonate | 3 |
| Vinyl acrylate | 3 |
| Ethyl 2-chlorobenzoate | 3 |
| 2,4-Dichlorophenyl 2,4-dichlorobenzoate [DCPDCB] | 3 |
| Propyl acetate | 3 |
| Phenyl propionate | 3 |
| 1-Naphthyl butyrate | 4 |
| Ethyl butyrate | 4 |
| Propylparaben | 4 |
| (−)-Methyl (*R*)-3-hydroxyvalerate | 4 |
| (+)-Methyl (*S*)-3-hydroxyvalerate | 4 |

| | |
|---|---|
| Benzylparaben | 4 |
| Propyl propionate | 4 |
| Methyl ferulate | 4 |
| Vinyl butyrate | 4 |
| 3-Methyl-3-buten-1-yl acetate | 4 |
| Ethyl 2-methylacetoacetate | 4 |
| Ethyl acetoacetate | 4 |
| Cyclohexyl butyrate | 4 |
| 2,4-Dichlorobenzyl 2,4-dichlorobenzoate [DCBDCB] | 4 |
| Butyl acetate | 4 |
| N-Benzyl-L-proline ethyl ester | 5 |
| N-Benzyl-D-proline ethyl ester | 5 |
| Ethyl (*R*)-(+)-4-chloro-3-hydroxybutyrate [E(R)CHB] | 5 |
| Ethyl (*S*)-(−)-4-chloro-3-hydroxybutyrate [E(S)CHB] | 5 |
| Benzoic acid 4-formyl-phenylmethyl ester [BFPME] | 5 |
| Butylparaben | 5 |
| Methyl hexanoate | 5 |
| Propyl butyrate | 5 |
| Isobutyl cinnamate | 5 |
| Ethyl 2-ethylacetoacetate | 5 |
| Ethyl propionylacetate | 5 |
| Hexyl acetate | 6 |
| Ethyl hexanoate | 6 |
| Phthalic acid diethyl ester | 6 |
| Benzyl (*R*)-(+)-2-hydroxy-3-phenylpropionate [BHPP] | 6 |
| Phenylethyl cinnamate | 6 |
| Geranyl acetate | 6 |
| Ethyl 3-oxohexanoate | 6 |
| n-Pentyl benzoate | 6 |
| Methyl octanoate | 7 |
| Propyl hexanoate | 7 |
| Diethyl-2,6-dimethyl 4-phenyl-1,4-dihydro pyridine-3,5-dicarboxylate [DDPDPDC] | 7 |
| Glyceryl triacetate | 8 |
| Octyl acetate | 8 |
| Ethyl octanoate | 8 |
| Methyl decanoate | 9 |
| (1*R*)-(-)-dimenthyl succinate | 9 |
| Ethyl decanoate | 10 |
| Glyceryl tripropionate | 11 |
| Methyl dodecanoate | 11 |
| Dodecanoyl acetate | 12 |
| Ethyl dodecanoate | 12 |
| Vinyl laurate | 12 |
| Methyl myristate | 13 |
| Glyceryl tributyrate | 14 |

| | |
|---|---|
| Ethyl myristate | 14 |
| Vinyl myristate | 14 |
| Pentadecyl acetate | 15 |
| Glucose pentaacetate | 15 |
| Methyl oleate | 16 |
| Vinyl palmitate | 16 |
| Vinyl oleate | 17 |
| Glyceryl trioctanoate | 26 |
| Triolein | 54 |

**Table S10: Distribution of $P_{EH}$ in $F_{EH}$ of the *experimental data set*.**

| **EH** | $F_{EH}$ [a] | $P_{EH}$ [b] |
|---|---|---|
| 026 | $F_I$ | 32 |
| 040 | | 27 |
| 041 | | 27 |
| 046 | | 23 |
| 071 | | 18 |
| 072 | | 18 |
| 075 | | 16 |
| 077 | | 16 |
| 090 | | 13 |
| 097 | | 11 |
| 101 | | 11 |
| 108 | | 9 |
| 110 | | 8 |
| 113 | | 8 |
| 115 | | 8 |
| 118 | | 6 |
| 131 | | 2 |
| 132 | | 2 |
| 142 | | 1 |
| 145 | | 1 |
| 051 | $F_{II}$ | 22 |
| 073 | | 17 |
| 088 | | 13 |
| 098 | | 11 |
| 102 | | 10 |
| 116 | | 7 |
| 136 | | 2 |
| 138 | | 2 |
| 140 | | 1 |
| 001 | $F_{IV}$ | 72 |
| 002 | | 71 |
| 003 | | 69 |
| 004 | | 67 |
| 005 | | 67 |
| 006 | | 66 |
| 008 | | 63 |
| 009 | | 61 |
| 010 | | 58 |
| 011 | | 53 |
| 012 | | 51 |
| 013 | | 49 |
| 014 | | 48 |
| 015 | | 42 |
| 016 | | 42 |
| 018 | | 38 |
| 021 | | 36 |
| 022 | | 35 |

| | | |
|---|---|---|
| 023 | | 34 |
| 025 | | 33 |
| 029 | | 31 |
| 035 | | 29 |
| 037 | | 28 |
| 039 | | 28 |
| 042 | | 27 |
| 043 | | 27 |
| 048 | | 23 |
| 052 | | 21 |
| 054 | | 21 |
| 067 | | 18 |
| 079 | | 14 |
| 086 | | 13 |
| 087 | | 13 |
| 091 | | 13 |
| 099 | | 11 |
| 119 | | 6 |
| 028 | | 31 |
| 031 | | 29 |
| 032 | | 29 |
| 045 | | 24 |
| 047 | | 23 |
| 049 | | 23 |
| 053 | | 21 |
| 055 | | 21 |
| 056 | | 21 |
| 057 | | 21 |
| 058 | | 21 |
| 065 | | 20 |
| 066 | | 19 |
| 068 | | 18 |
| 074 | | 17 |
| 076 | | 16 |
| 081 | *F*V | 14 |
| 082 | | 14 |
| 083 | | 14 |
| 092 | | 12 |
| 094 | | 12 |
| 096 | | 11 |
| 010 | | 11 |
| 103 | | 10 |
| 104 | | 10 |
| 107 | | 9 |
| 109 | | 9 |
| 111 | | 8 |
| 114 | | 8 |
| 120 | | 5 |
| 123 | | 4 |
| 127 | | 4 |
| 128 | | 4 |

| | | |
|---|---|---|
| 030 | | 30 |
| 034 | | 29 |
| 059 | FVI | 21 |
| 061 | | 20 |
| 085 | | 13 |
| 020 | | 37 |
| 064 | | 20 |
| 084 | | 13 |
| 093 | *F*VII | 12 |
| 112 | | 8 |
| 139 | | 2 |
| 007 | | 64 |
| 024 | | 34 |
| 027 | | 32 |
| 069 | | 18 |
| 078 | | 15 |
| 089 | FVIII (serine beta-lactamase like) | 13 |
| 095 | | 11 |
| 124 | | 4 |
| 133 | | 2 |
| 141 | | 1 |
| 129 | | 3 |
| 044 | | 25 |
| 070 | | 18 |
| 126 | CE (carbohydrate esterase like) | 4 |
| 134 | | 2 |
| 135 | | 2 |
| 137 | | 2 |
| 017 | | 39 |
| 019 | | 37 |
| 033 | | 29 |
| 036 | | 28 |
| 038 | C-C MCPh | 28 |
| 050 | | 22 |
| 060 | | 21 |
| 062 | | 20 |
| 063 | | 20 |
| 105 | Cyclase-like esterase | 10 |
| CalB | Yeast class | 68 |
| CalA | | 36 |
| 080 | | 14 |
| 106 | | 9 |
| 117 | | 6 |
| 121 | | 5 |
| 122 | Unclassified | 5 |
| 125 | | 4 |
| 130 | | 2 |
| 144 | | 1 |
| 143 | | 1 |

[a] EH families based on the Arpigny and Jaeger classification(35) **(see section 2.1)**.

[b] Experimentally measured substrate promiscuity level of EHs provided by Martínez-Martínez *et al.* (1) **(see section 2.1)**.

**Table S11: $P_{EH}$ and $Vol_{eff}$ of comparative models of EHs of the *flexibility data set* with known crystal structures.**

| EH | $P_{EH}$[a] | $Vol_{eff}$ [Å$^3$][b] |
|---|---|---|
| 001 | 72 | 166.667 |
| CalB | 68 | 200.000 |
| CalA | 36 | 1000.000 |
| 023 | 34 | 90.909 |
| 037 | 28 | 166.667 |
| 060 | 21 | 250.000 |
| 096 | 11 | 34.483 |
| 102 | 10 | 38.462 |
| 105 | 10 | n.d.[c] |
| 107 | 9 | 28.571 |
| 115 | 8 | 71.429 |

[a] Experimentally determined substrate promiscuity level of EHs provided by Martínez-Martínez *et al.* (1) **(see section 2.1)**.

[b] Computed active site effective volumes of EHs provided by Martínez-Martínez *et al.* (1) **(see section 2.1)**.

[c] Not determined.

**Table S12: $P_{EH}$ and $Vol_{eff}$ of comparative models of EHs of the *flexibility data set* without known crystal structures.**

| EH | $P_{EH}$[a] | $Vol_{eff}$ [Å$^3$][b] |
|---|---|---|
| 005 | 67 | 200.000 |
| 010 | 58 | 200.000 |
| 011 | 53 | 83.333 |
| 012 | 51 | 333.333 |
| 013 | 49 | 333.333 |
| 014 | 48 | 200.000 |
| 015 | 42 | 166.667 |
| 016 | 42 | 333.333 |
| 029 | 31 | 500.000 |
| 030 | 30 | 66.667 |
| 033 | 29 | 166.667 |
| 034 | 29 | 32.258 |
| 042 | 27 | 200.000 |
| 043 | 27 | 66.667 |
| 048 | 23 | 111.111 |
| 061 | 20 | 111.111 |
| 070 | 18 | 43.478 |
| 074 | 17 | 58.824 |
| 083 | 14 | 58.824 |
| 092 | 12 | 41.667 |
| 093 | 12 | 37.037 |
| 094 | 12 | 45.455 |
| 125 | 4 | 19.231 |
| 127 | 4 | 55.556 |

[a] Experimentally determined substrate promiscuity level of EHs provided by Martínez-Martínez *et al.* (1) **(see section 2.1)**.

[b] Computed active site effective volumes of EHs provided by Martínez-Martínez *et al.* (1) **(see section 2.1)**.

[c] Not determined.

### Supplemental Figures

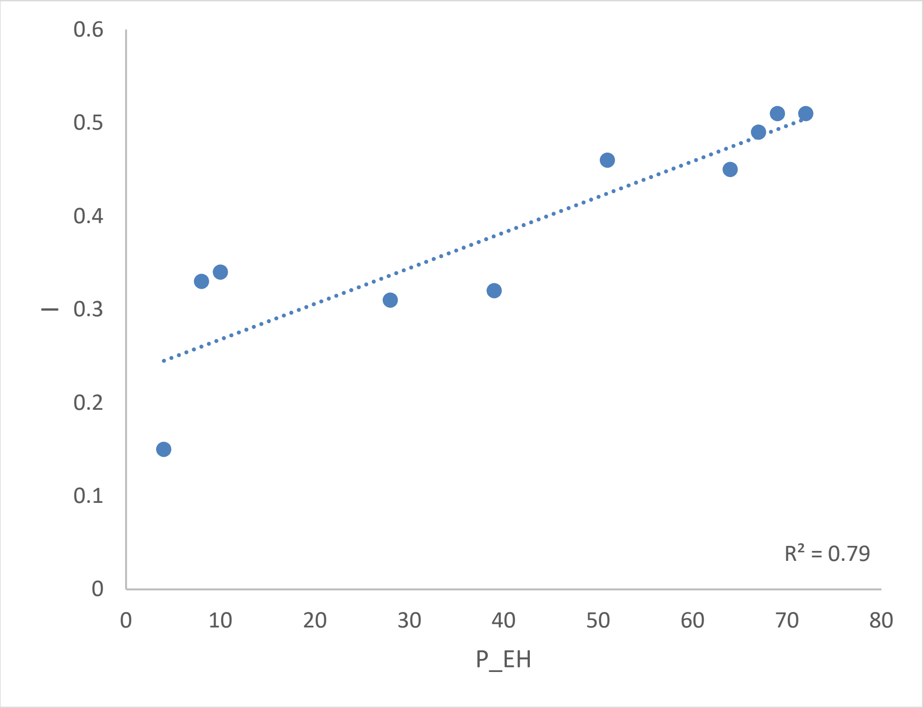


**Figure S1: Correlation of the promiscuity index *I* (Eq. S4) versus $P_{EH}$ for the ten EHs for which $k_{cat}$ and $K_m$ values were determined.**

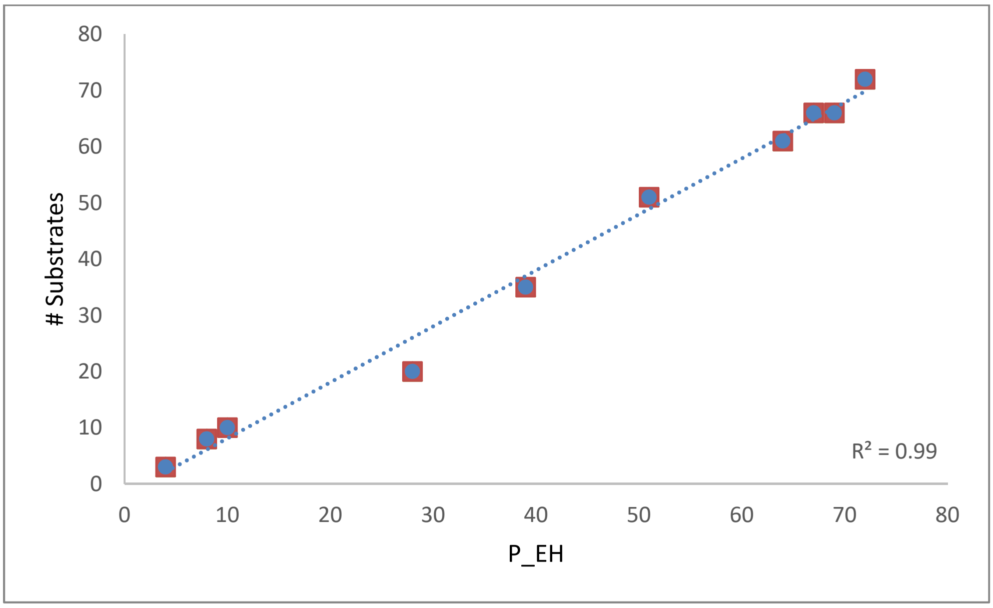


**Figure S2: Correlation of the number of substrates with $k_{cat}$ / $K_m$ > 0.05 $min^{-1}$ $M^{-1}$ versus $P_{EH}$ for the ten EHs for which $k_{cat}$ and $K_m$ values were determined.**

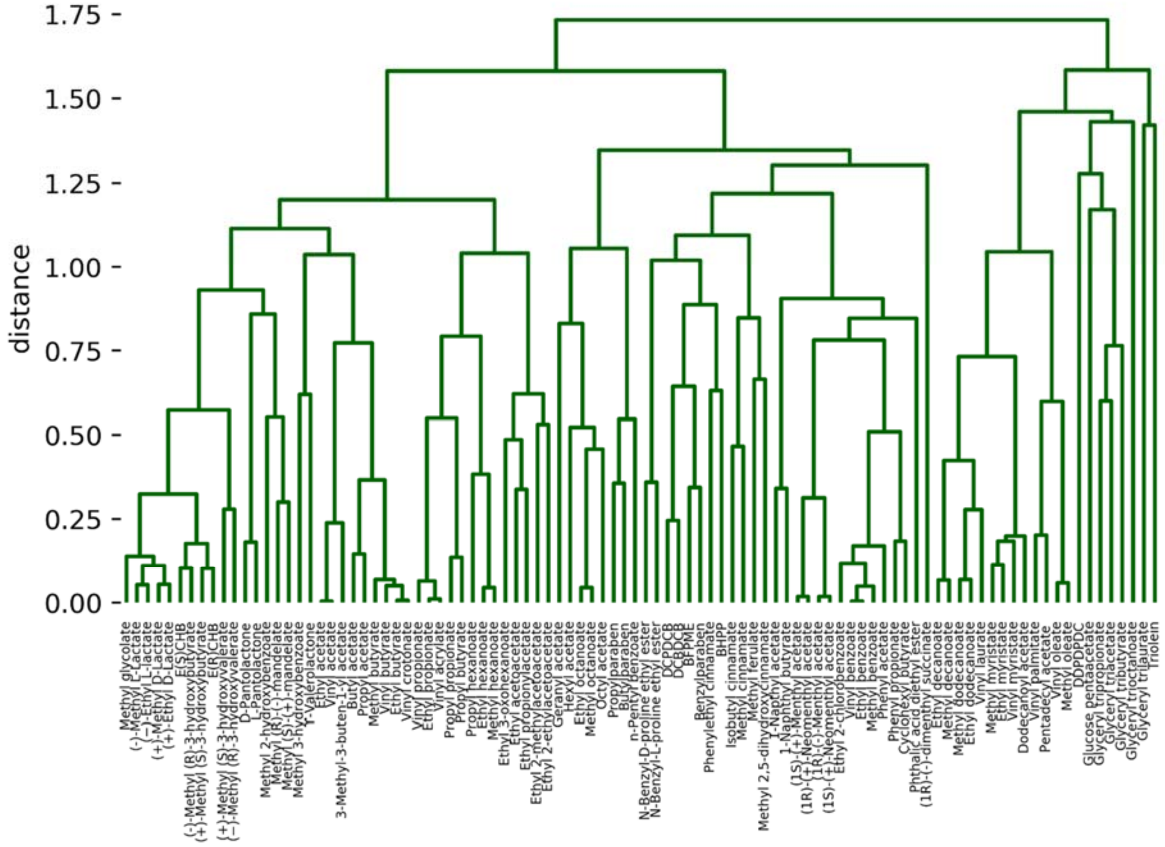


**Figure S3: Hierarchical clustering of ester substrates.** The complete linkage clustering method was used to define clusters using the all-*vs*.-all pairwise distance matrix calculated for all 96 compounds from the respective matrix of maximum pairwise Tanimoto-Combo similarity scores $\delta_{ij}$.

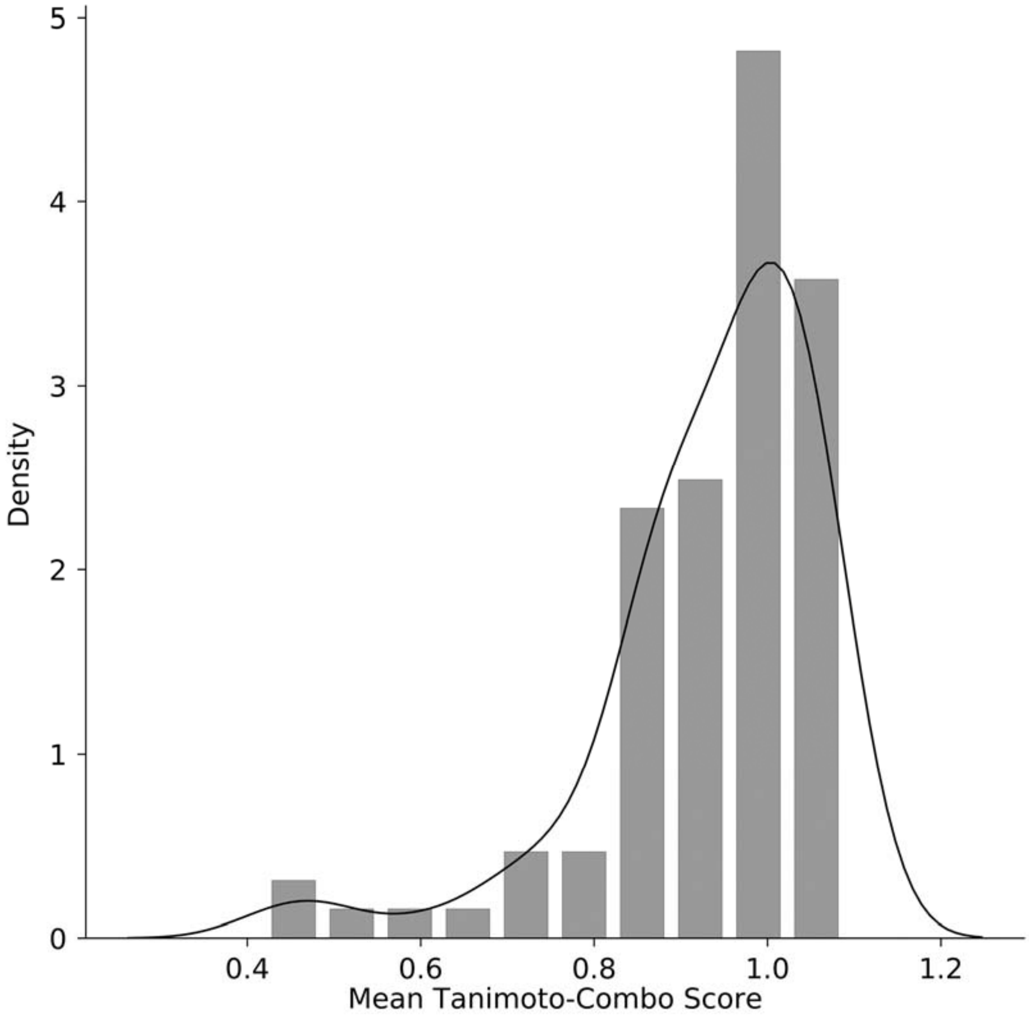


**Figure S4: Distribution of the mean Tanimoto-Combo similarity scores $\delta_i$.** The mean scores were calculated for each of the 96 ester substrates as the respective row-average in the all-*vs*.-all matrix of maximum pairwise Tanimoto-Combo similarity scores $\delta_{ij}$.

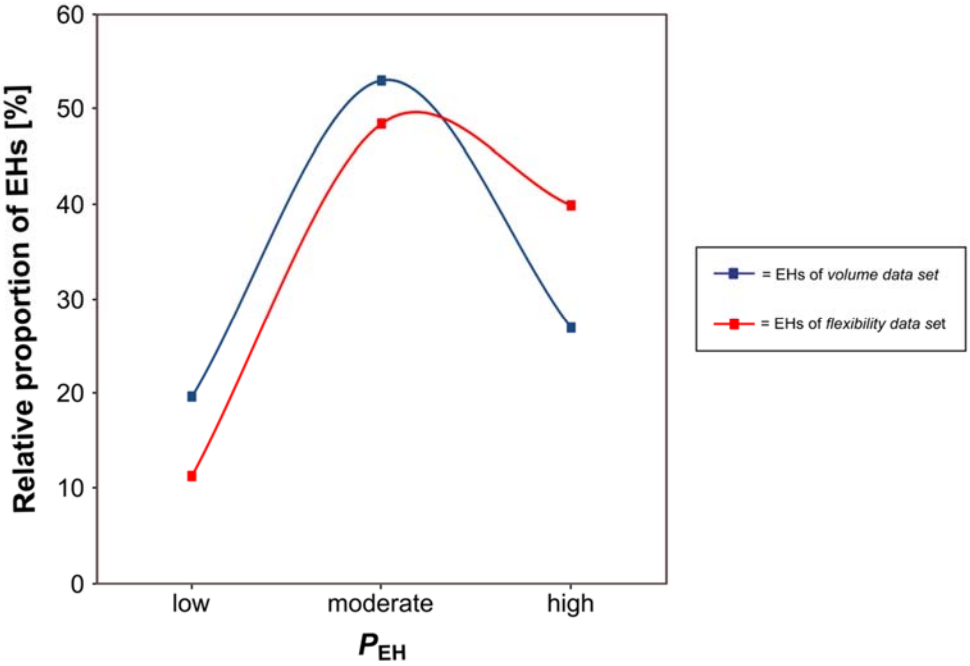


**Figure S5: Comparison between the *volume data set* and the *flexibility data set* regarding $P_{EH}$.** Relative proportions of EHs constituting the *volume data set* (red line) and the *flexibility data set* (blue line) regarding $P_{EH}$ determined with a kinetic pH indicator assay (2-4) by Martínez-Martínez *et al.* (1). $P_{EH}$ is defined as *low* if the EH hydrolyzes ≤ 9 esters, as *moderate* if the EH hydrolyzes between 10 and 29 esters, and as *high* if the EH hydrolyzes ≥ 30 esters.

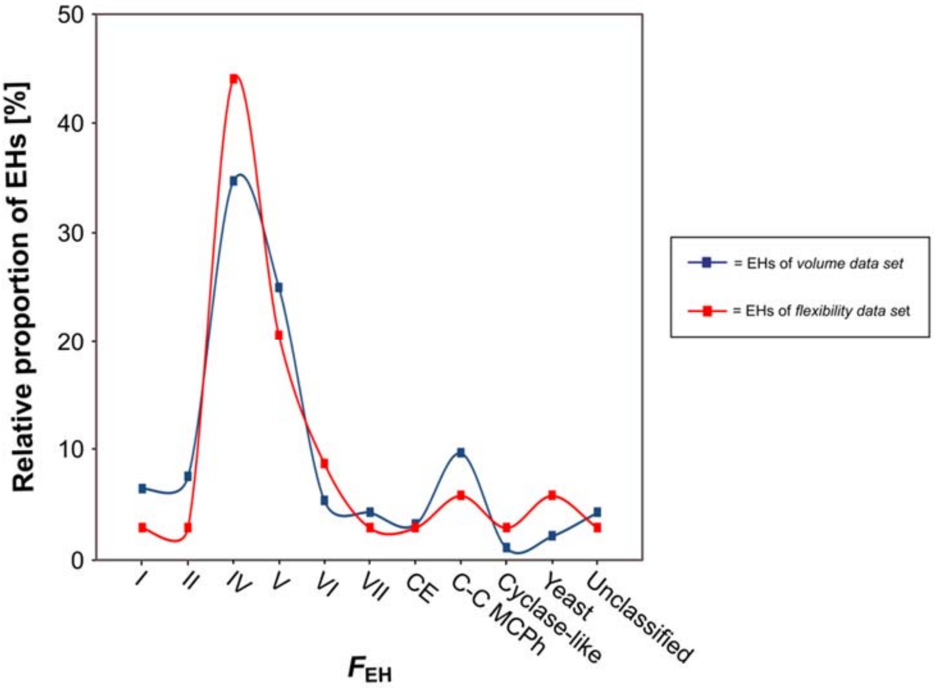


**Figure S6: Comparison between the *volume data set* and the *flexibility data set* regarding $F_{EH}$.** Relative proportions of EHs constituting the *volume data set* (red line) and the *flexibility data set* (blue line) regarding $F_{EH}$ based on the Arpigny and Jaeger classification (35).

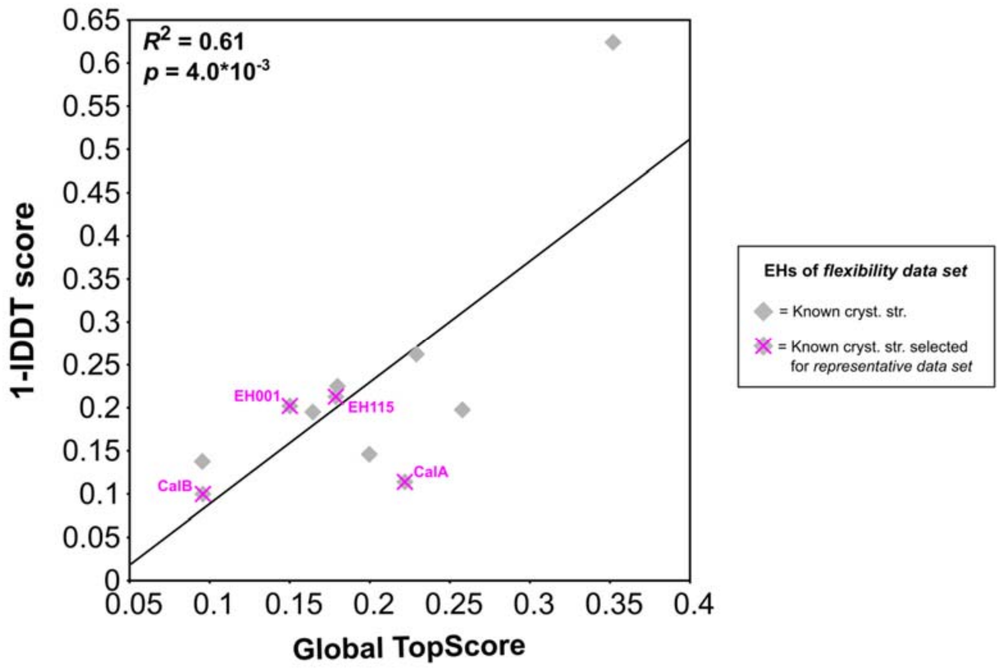


**Figure S7: TopScore performance on comparative models of EHs of the *flexibility data set* with known crystal structures.** Correlation between 1 - lDDT scores and global TopScores for comparative models of EHs of the *flexibility data set* with known crystal structures. The comparative models were generated by TopModel (36) (excluding the known crystal structures as templates) and evaluated by TopScore (15). The lDDT scores were computed by the lDDT web server from Swiss-Model (16) from comparisons of the comparative models of these EHs against the known crystal structures as experimental references. The EHs belonging to the *representative data set* are indicated by magenta crosses.

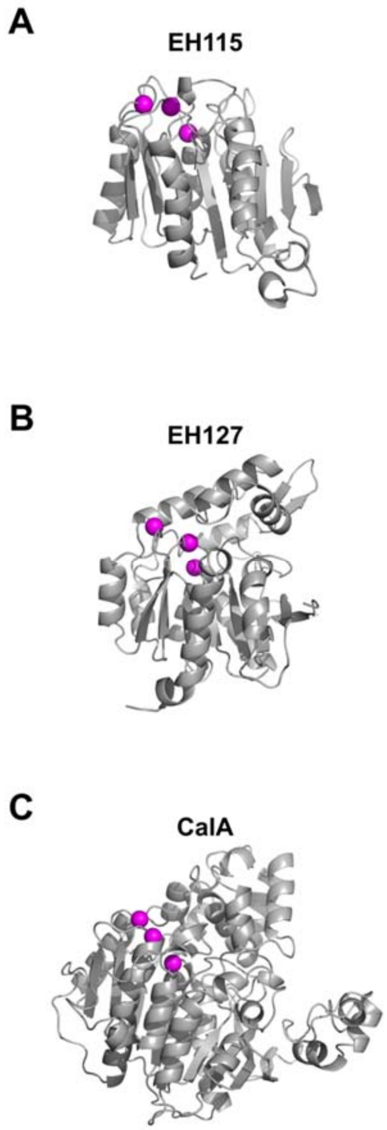


**Figure S8: Substrate-accessibility of EHs of the *representative data set*.** CAVER results (37) of comparative models of **(A)** EHs with known crystal structures and lowest (EH115) or highest $P_{EH}$ (EH001), **(B)** EHs with unknown crystal structures and lowest (EH127) or highest $P_{EH}$ (EH005), and **(C)** commercial EHs with lowest (CalA) or highest $P_{EH}$ (CalB). CARs (magenta spheres) are either located on the protein surface or are buried and connected with the surface by tunnels (blue, green, and red spheres).

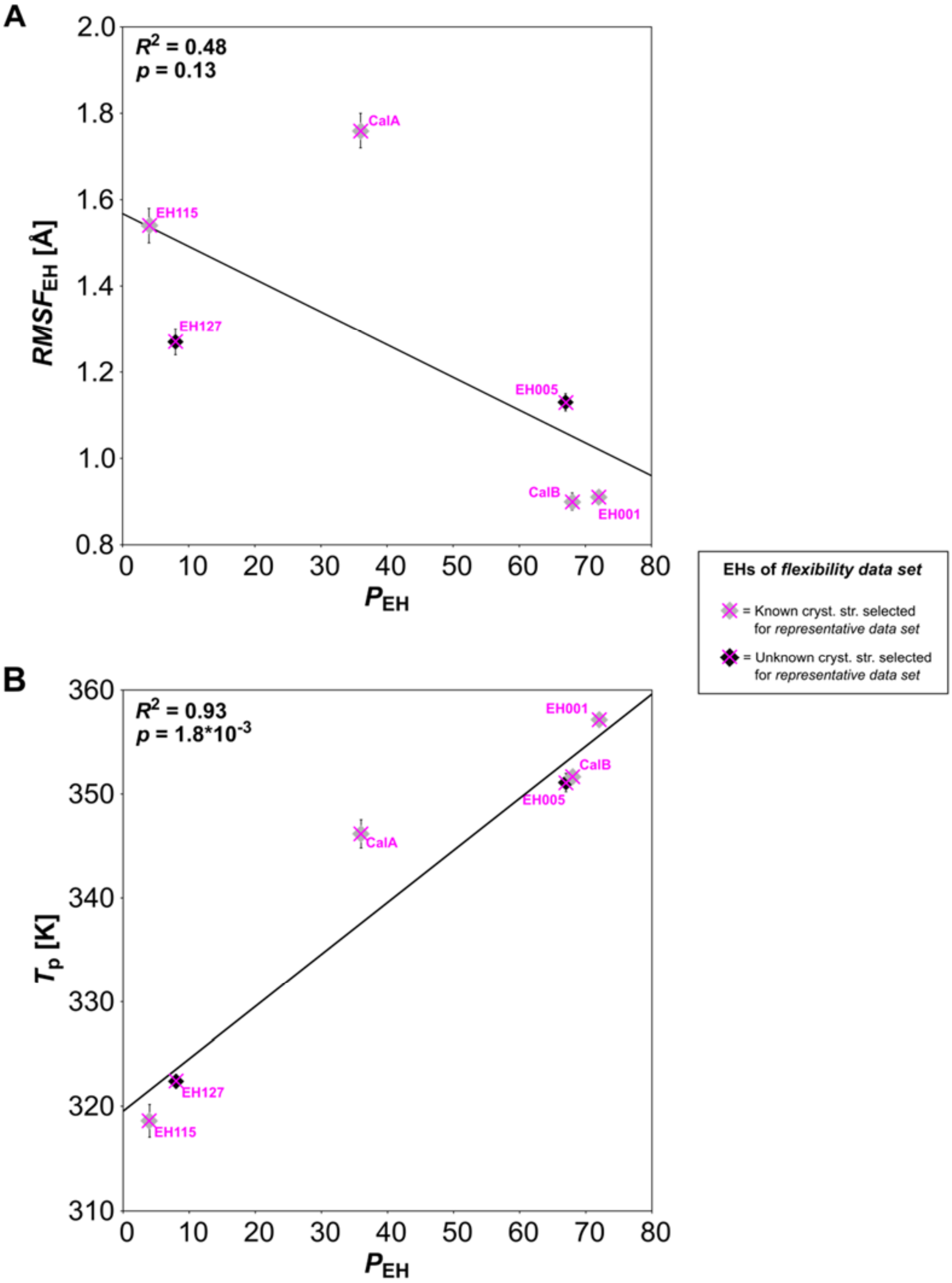


**Figure S9: Correlation of *RMSF*$_{EH}$ or *T*$_p$ *versus P*$_{EH}$ of the *representative data set*. (A)** Correlation between *RMSF*$_{EH}$ based on the MD trajectories and *P*$_{EH}$ of the *representative data set*. **(B)** Correlation between $T_p$ based on the *global* index $H_{type2}$ (25) computed by CNA and $P_{EH}$ of the *representative data set*. Data points colored grey (black) and indicated by magenta crosses represent comparative models of EHs with (un)known crystal structures. Error bars show the SEM over five independent MD simulations of 1 µs length each.

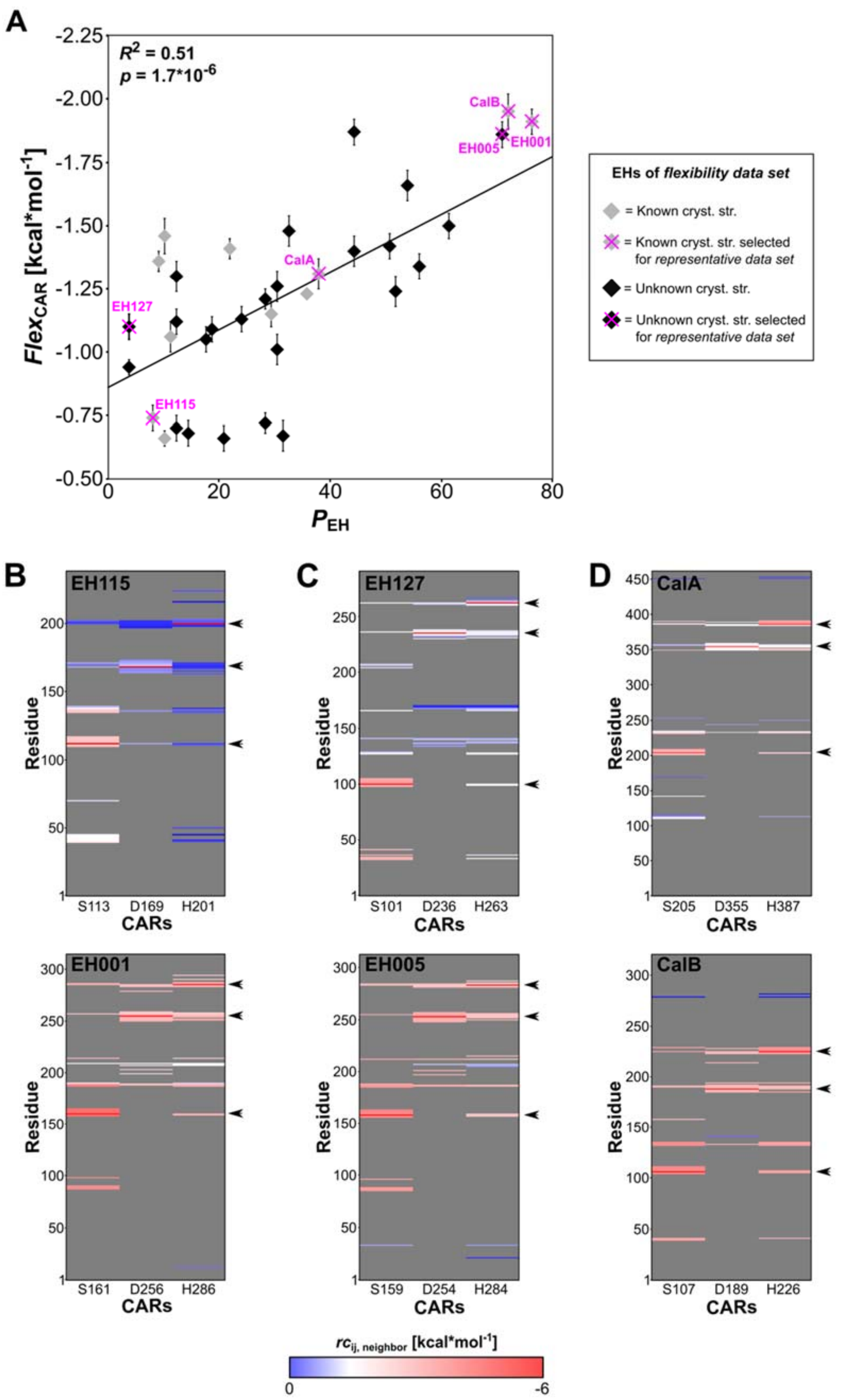


**Figure S10: Correlation of *Flex*$_{CAR}$ *versus* *P*$_{EH}$. (A)** Correlation between predicted $Flex_{CAR}$ based on the local index $rc_{ij;neighbor}$ and $P_{EH}$ for the *flexibility data set*. Data points colored grey (black) represent homology models of EHs with (un)known crystal structures. The *representative data set* is indicated by magenta crosses. Error bars show the SEM over five independent MD simulations of 1 μs length each.. $rc_{ij;neighbor}$ of CARs of **(B)** EHs with known crystal structures and lowest (EH115) or highest $P_{EH}$ (EH001), **(C)** EHs with unknown crystal structures and lowest (EH127) or highest $P_{EH}$ (EH005), and **(D)** commercial EHs with lowest (CalA) or highest $P_{EH}$ (CalB). A red (blue) color indicates that a rigid contact between CARs and other residues within 5 Å distance is more (less) stable (see color scale at the bottom). The rigid contacts for all other residue pairs are colored grey. Black arrow heads indicate positions of CARs.

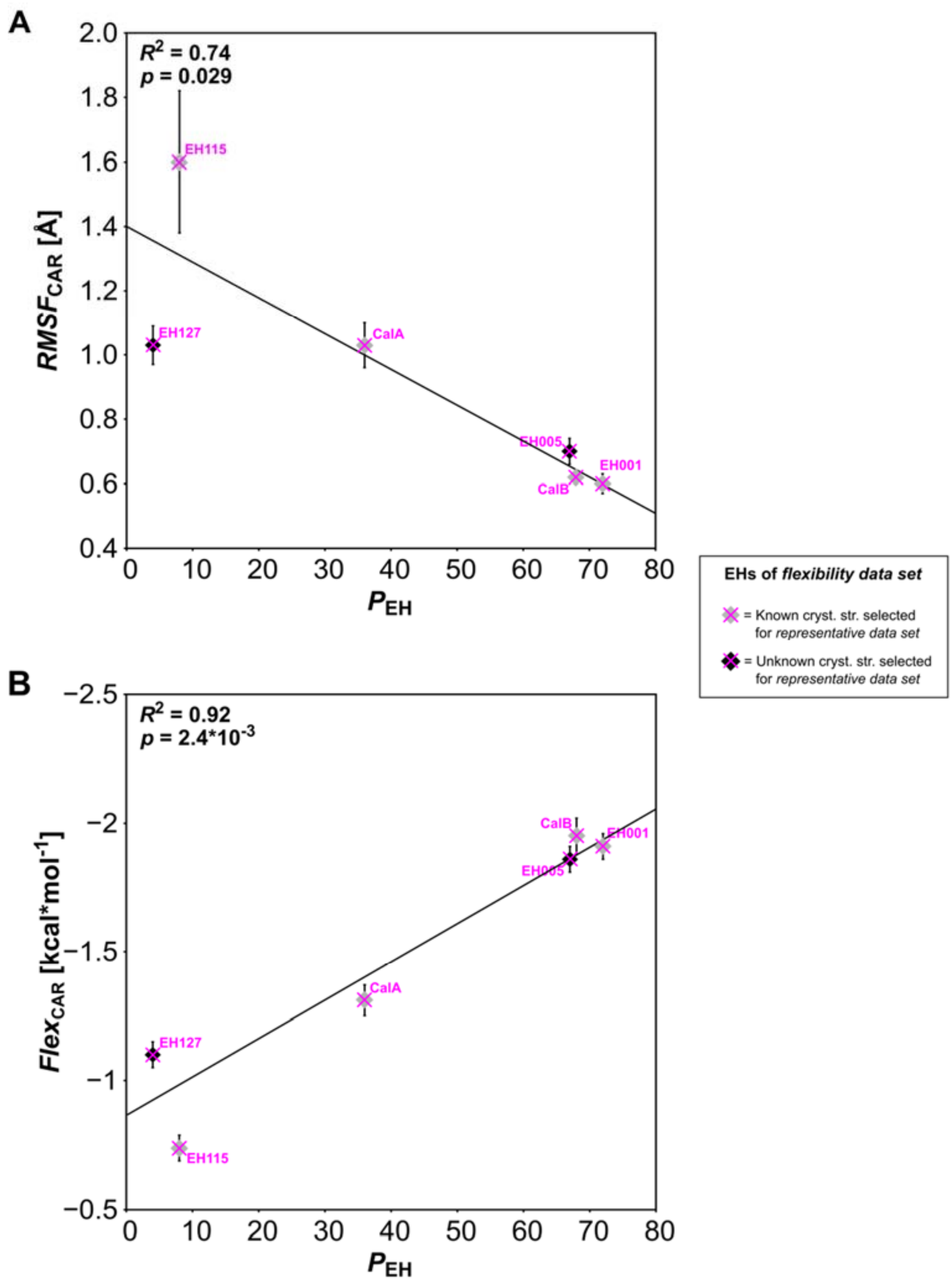


**Figure S11: Correlation of *RMSF*$_{CAR}$ or *Flex*$_{CAR}$ *versus P*$_{EH}$ of the *representative data set*.** **(A)** Correlation between *RMSF*$_{CAR}$ based on the MD trajectories and *P*$_{EH}$ of the *representative data set*. **(B)** Correlation between *Flex*$_{CAR}$ based on CNA and *P*$_{EH}$ of the *representative data set*. Data points colored grey (black) and indicated by magenta crosses represent comparative models of EHs with (un)known crystal structures. Error bars show the SEM over five independent MD simulations of 1 µs length each.

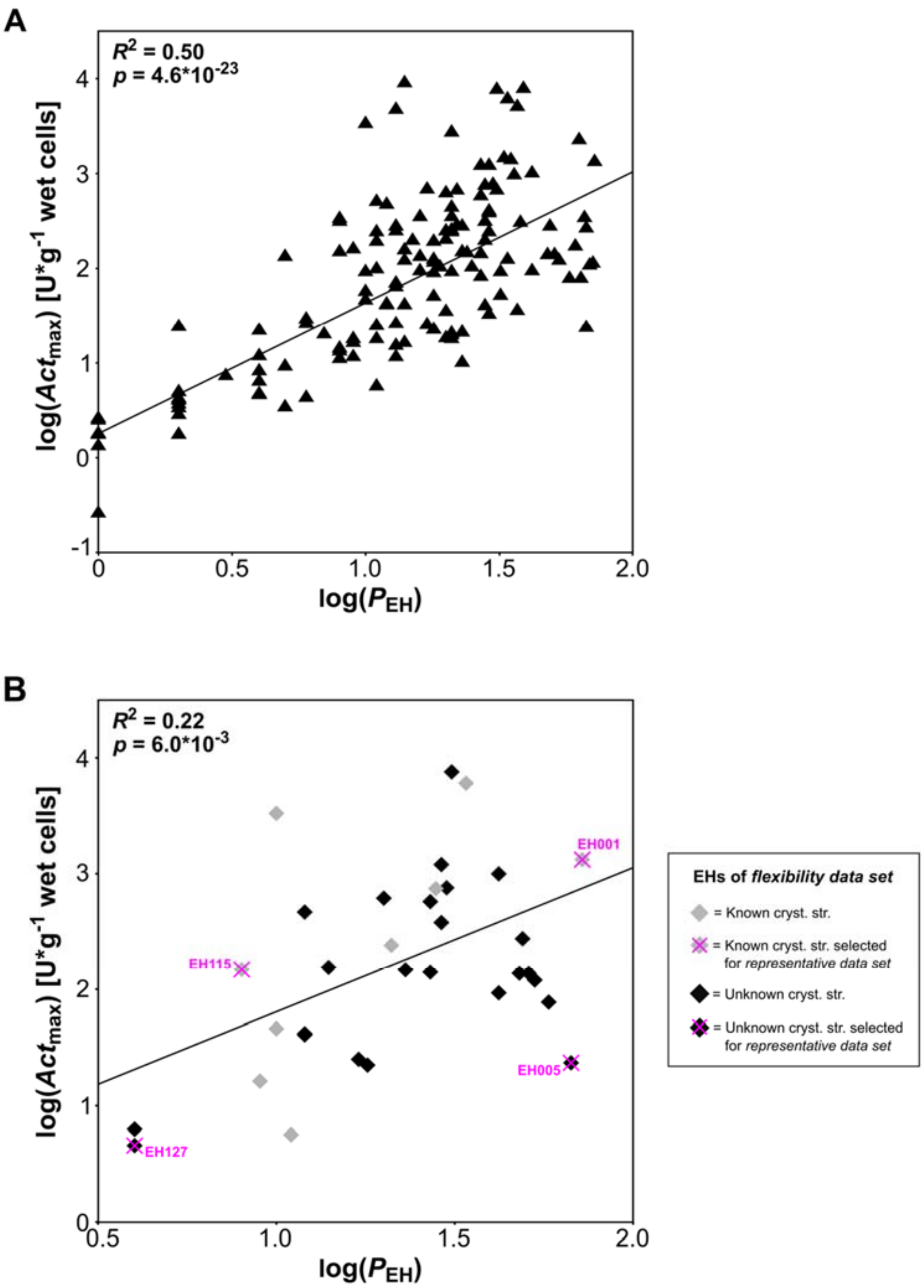


**Figure S12: Correlation of log($Act_{max}$) *versus* log($P_{EH}$).** Correlation between log($Act_{max}$) and log($P_{EH}$) for **(A)** the *experimental data set* and **(B)** the *flexibility data set* containing EHs with known crystal structures (grey data points), EHs with unknown crystal structures (black data points), and EHs constituting the *representative data set* (magenta crosses). The EHs were screened against 96 different esters in a kinetic pH indicator assay (2-4) that provided $Act_{max}$ given in U (g wet cells)$^{-1}$. CalA and CalB preparations were excluded because $Act_{max}$ was given in U (g total protein)$^{-1}$. The assays were performed as triplicates with STD ≤ 1%.

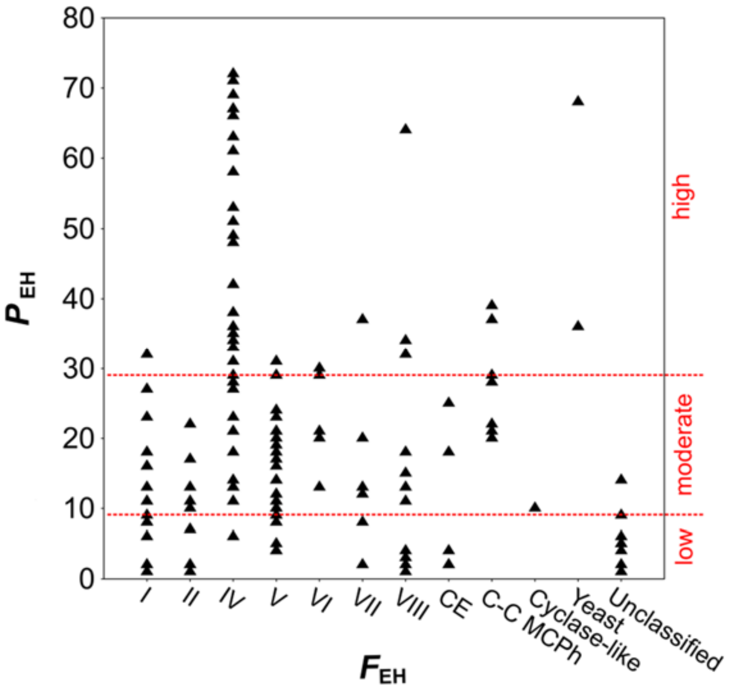


**Figure S13: Distribution of $P_{EH}$ in $F_{EH}$ of the *experimental data set*.** Distribution of $P_{EH}$ determined with a kinetic pH indicator assay (2-4) by Martínez-Martínez *et al.* (1) in $F_{EH}$ based on the Arpigny and Jaeger classification (35) of the *experimental data set*. $P_{EH}$ is defined as *low* if the EH hydrolyzes ≤ 9 esters, as *moderate* if the EH hydrolyzes between 10 and 29 esters, and as *high* if the EH hydrolyzes ≥ 30 esters.

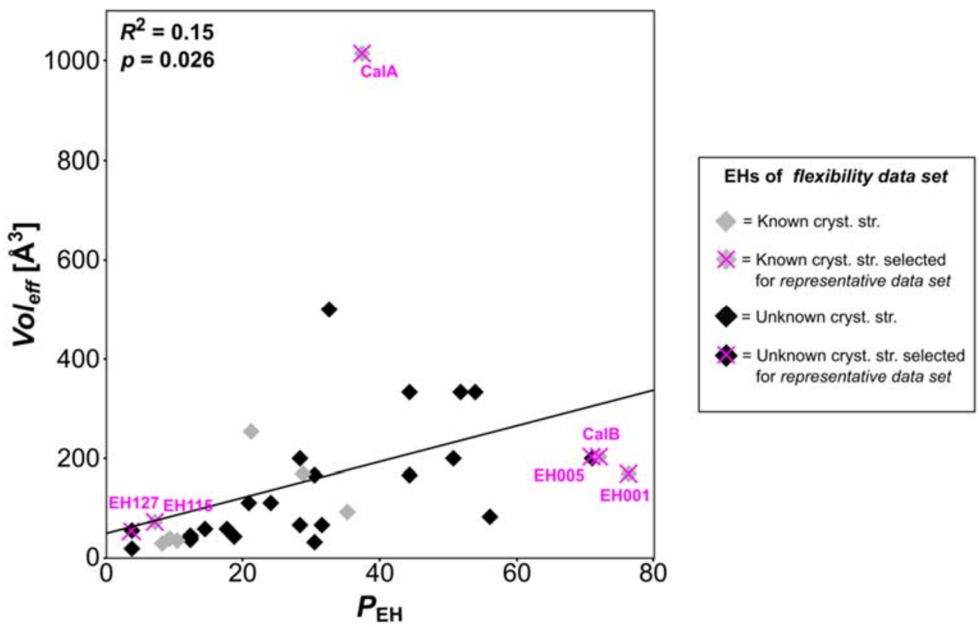


**Figure S14: Correlation of *$Vol_{eff}$ versus* $P_{EH}$.** Correlation between *$Vol_{eff}$* and $P_{EH}$ of the *flexibility data set*. *$Vol_{eff}$* represents the topology of the catalytic environment in terms of the active site cavity volume (*$Vol_{cav}$*) computed by Fpocket (5) per relative solvent-accessible surface area (*$SASA_{rel}$*) computed by GetArea webserver (6). Data points colored grey (black) represent comparative models of EHs with (un)known crystal structures. The *representative data set* is indicated by magenta crosses.
